# Nuclear deformation causes DNA damage by increasing replication stress

**DOI:** 10.1101/2020.06.12.148890

**Authors:** Pragya Shah, Chad M. Hobson, Svea Cheng, Marshall Colville, Matthew Paszek, Richard Superfine, Jan Lammerding

## Abstract

Cancer metastasis, i.e., the spreading of tumor cells from the primary tumor to distant organs, is responsible for the vast majority of cancer deaths. In the process, cancer cells migrate through narrow interstitial spaces substantially smaller in cross-section than the cell. During such confined migration, cancer cells experience extensive nuclear deformation, nuclear envelope rupture, and DNA damage. The molecular mechanisms responsible for the confined migration-induced DNA damage remain incompletely understood. While in some cell lines, DNA damage is closely associated with nuclear envelope rupture, we show that in others, mechanical deformation of the nucleus is sufficient to cause DNA damage, even in the absence of nuclear envelope rupture. This deformation-induced DNA damage, unlike nuclear envelope rupture-induced DNA damage, occurs primarily in S/G2 phase of the cell cycle and is associated with replication forks. Nuclear deformation, resulting from either confined migration or external cell compression, increases replication stress, possibly by increasing replication fork stalling, providing a molecular mechanism for the deformation-induced DNA damage. Thus, we have uncovered a new mechanism for mechanically induced DNA damage, linking mechanical deformation of the nucleus to DNA replication stress. This mechanically induced DNA damage could not only increase genomic instability in metastasizing cancer cells, but could also cause DNA damage in non-migrating cells and tissues that experience mechanical compression during development, thereby contributing to tumorigenesis and DNA damage response activation.

## Introduction

Cell migration is important for various developmental processes and immune surveillance [1, 2]. In addition, cell migration is essential for tumor cell invasion and metastasis, which is responsible for more than 80% of all cancer deaths [3]. During metastasis, cancer cells disseminate from the primary tumor, invade through the surrounding extracellular matrix and into neighboring tissues, ultimately spreading to distant organs via the circulation and lymphatic system [3]. In this process, cancer cells encounter interstitial spaces of the order of 0.1-20 μm in diameter, i.e., smaller than the size of the cell nucleus [4–6]. Migration through such confined environments puts considerable mechanical stress on the cell nucleus, which constitutes the largest and stiffest organelle [7, 8]. As a result, cells frequently experience severe nuclear deformation and nuclear envelope (NE) rupture during confined migration [9–16]. Transient loss of NE integrity allows uncontrolled exchange between the nucleoplasm and cytoplasm, exposes the genomic DNA to cytoplasmic components such as nucleases, and leads to DNA damage [9–11, 13, 15, 17–23]. Although cells rapidly repair their NE and continue to survive and migrate [10, 11, 13], the acquired DNA damage can increase genomic instability in these cancer cells [9, 14, 15, 19, 23, 24] which could further enhance their metastatic potential and resistance to therapies.

While it is now well recognized that confined migration can cause DNA damage, the underlying molecular mechanism remains incompletely understood. Recent reports have implicated loss of DNA repair factors during NE rupture or local exclusion of repair factors due to nuclear deformation as possible mechanisms [15–19]. Exposure to cytoplasmic DNases such as TREX1 following NE rupture [21–23] and mislocalization of organelles like mitochondria post NE rupture [25] have also been suggested as cause of DNA damage. Contributing to the uncertainty about the molecular mechanism responsible for the confined migration-induced DNA damage is that previous studies have provided at times conflicting results on the association between DNA damage and NE rupture. While some studies reported that DNA damage requires NE rupture [13–15, 26, 27], others found that DNA damage can occur in the absence of NE rupture as cells squeeze their nuclei through tight spaces [10, 28].

Using time-lapse microscopy and a broad panel of cell lines co-expressing fluorescent reporters for DNA damage and NE rupture, we found that in some cell lines, mechanical deformation of the nucleus is sufficient to cause DNA damage, while in other cell lines DNA damage is primarily associated with NE rupture. These results provide an explanation for the varied results in previous studies. Furthermore, we show that nuclear deformation-induced DNA damage frequently occurred at replication forks, and that nuclear deformation during confined migration or external compression led to increased replication stress. Thus, we demonstrate a novel mechanism by which deformation of the nucleus can cause DNA damage in the absence of NE rupture. Intriguingly, deformation-induced DNA damage does not require cell migration, but is also seen in stationary cells subjected to physical compression. Thus, this mechanism could have broad implications during development and in tissues subjected to regular compression, such as solid tumors, skin, or cartilage, where the DNA damage could promote genomic instability and activate apoptosis or senescence pathways.

## Results

### NE rupture and nuclear deformation lead to DNA damage during confined migration

To address the specific cause of DNA damage during confined migration, we performed a systematic study using a panel of cells consisting of two breast cancer cell lines (MDA-MB-231 and BT-549), a fibrosarcoma cell line (HT1080), and two normal human cell lines (RPE-1 and human skin fibroblasts) that had previously been reported to exhibit DNA damage during confined migration [10, 13]. Cells were modified to stably express a fluorescent reporter for DNA damage, 53BP1-mCherry, which localizes to DNA double strand breaks (DSBs) [10, 13, 29]. Treatment with Phleomycin, a DSB inducing agent [30], and staining for ɤ-H2AX, which accumulates at DSBs, was used to validate the 53BP1-mCherry DNA damage reporter (Suppl. Fig. S1A-C). To detect NE rupture, cells were modified to co-express a NE rupture reporter, consisting of a green fluorescent protein with a nuclear localization sequence (NLS-GFP), which spills from the nucleus into the cytoplasm upon NE rupture and is re-imported into the nucleus once NE integrity has been restored [10, 11]. Using time-lapse microscopy, we monitored cells for DNA damage, NE rupture, and nuclear deformation as they migrated through collagen matrices or custom-built microfluidic devices that mimic the interstitial spaces found in *vivo* [10, 11, 31, 32]. The microfluidic devices contain channels with constrictions either 1 × 5 μm^2^ or 2 × 5 μm^2^ in cross-section that require extensive nuclear deformation, and larger control channels with 15 × 5 μm^2^ openings that do not require substantial nuclear deformation while still providing a 3-D cell environment.

For all cell types, migration of cells through the ≤ 2 × 5 μm^2^ constrictions led to a higher increase in DNA damage than migration through the 15 × 5 μm^2^ control channels, as seen by comparing the number of 53BP1-mCherry foci in individual cells before, during, and after passage through a constriction (Fig. 1A-B, 1C-D, Suppl. Fig. S1D-F). In HT1080 fibrosarcoma cells, RPE-1 retinal epithelial cells, and immortalized human skin fibroblasts, the DNA damage occurred predominantly following NE rupture (Fig. 1A, 1C, 1G; Suppl. Fig. S1G-H, S1J; Suppl. Video 1), consistent with a previous report in RPE-1 cells [13]. In the MDA-MB-231 and BT-549 breast cancer cells, in contrast, DNA damage predominantly occurred in the absence of NE rupture as the cell squeezed the nucleus through the tight constrictions (Fig. 1D-G; Suppl. Fig. S1I-J; Suppl. Video 2). While NE rupture also led to an increase in DNA damage in these cells, the extent of damage was much lower when compared to damage induced by nuclear deformation in the absence of rupture. These data suggest that while DNA damage is associated with NE rupture in some cell lines, nuclear deformation is sufficient to induce DNA damage, even without NE rupture, in other cell lines. Furthermore, the cell line-specific differences may explain the conflicting findings obtained in previous studies. Collectively, our results indicate that DNA damage during confined migration can result from two distinct, albeit overlapping events: nuclear deformation, as primarily seen in MDA-MB-231 and BT-549 breast cancer cells; and NE rupture (combined with nuclear deformation), as seen in HT1080, human fibroblasts and RPE-1 cells. To test whether the differences among the cell lines in the cause of DNA damage during confined migration could be attributed to variability in their nuclear deformability, we compared the nuclear elastic modulus and levels of lamins A/C, which are major contributors of nuclear deformability [33, 34], between these different cell lines. As expected, we observed a correlation between resistance to stretching of the nuclear surface and levels of lamin A/C within individual cell lines (Suppl. Fig. S2B-C); however neither nuclear deformability, lamin A/C levels, nor migration speed through the ≤ 2 × 5 μm^2^ constrictions revealed any consistent correlation with the cause of DNA damage (Suppl. Fig. S2), suggesting that other mechanisms are at play.

**Figure 1:**
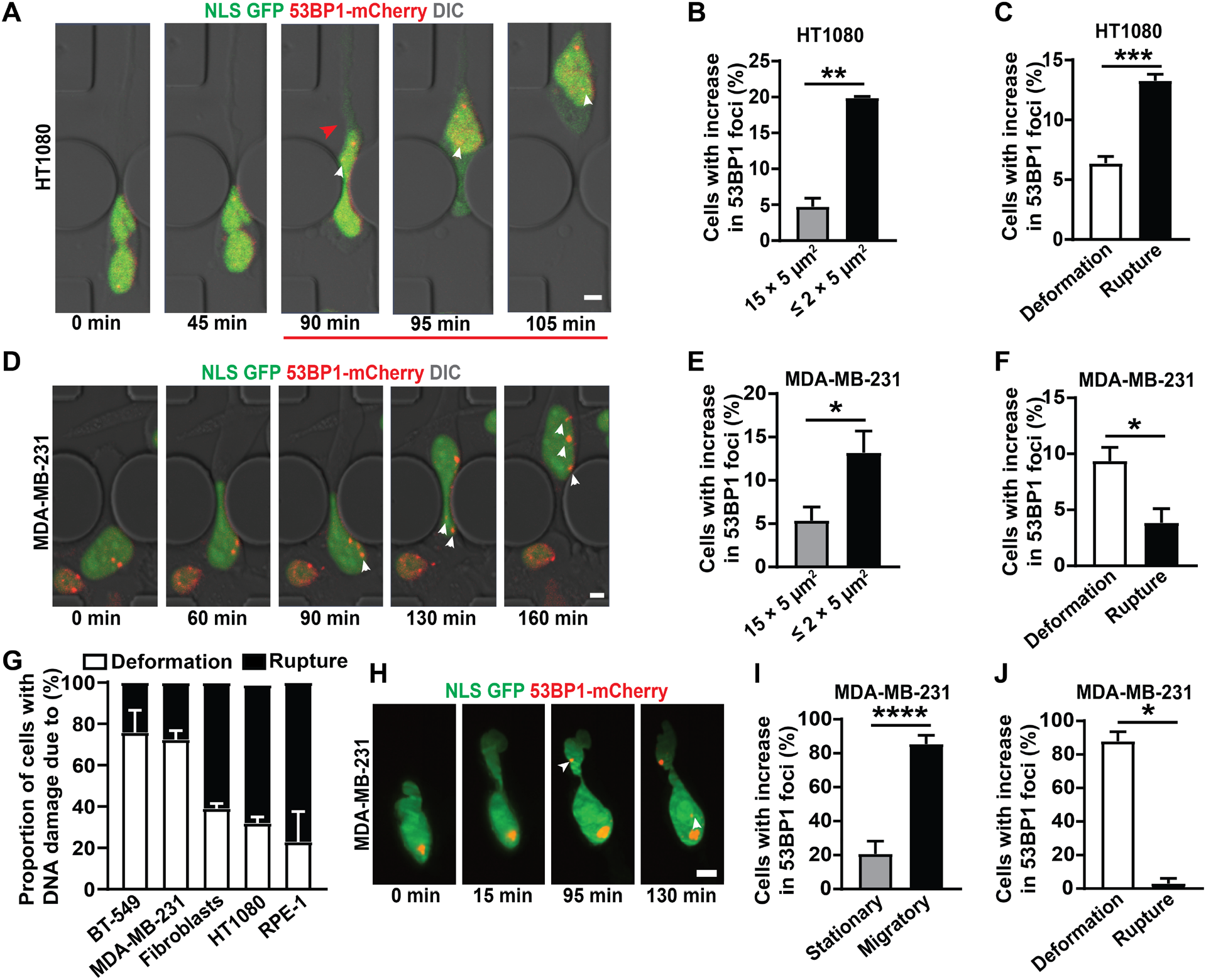
NE rupture and nuclear deformation lead to DNA damage during confined migration. **(A)** Representative image panel showing a HT1080 fibrosarcoma cell co-expressing NLS-GFP and 53BP1-mCherry exhibiting DNA damage following NE rupture during migration through a 1 × 5 μm^2^ constriction in the microfluidic device. Red arrowhead indicates start of NE rupture; the red line indicates the duration of NE rupture; white arrowheads indicate newly occurring 53BP1-mCherry foci. Scale bar: 5 μm **(B)** Percentage of HT1080 cells with new DNA damage (53BP1-mCherry foci) during migration through small (≤ 2 × 5 μm^2^) constrictions (*n* = 372 cells) or 15 × 5 μm^2^ control channels (*n* = 268 cells). **, *p* < 0.01 based on unpaired *t*-test with Welch’s correction. **(C)** Percentage of HT1080 cells in which new DNA damage during migration through ≤ 2 × 5 μm^2^ constrictions was associated with either NE rupture or with nuclear deformation in the absence of NE rupture. *n* = 372 cells; ***, *p* < 0.001 based on unpaired *t*-test with Welch’s correction. **(D)** Representative image sequence showing a MDA-MB-231 breast cancer cell co-expressing NLS-GFP and 53BP1-mCherry experiencing new DNA damage during migration through a 2 × 5 μm^2^ constriction. White arrowheads indicate newly occurring 53BP1-mCherry foci. Scale bar: 5 μm **(E)** Percentage of MDA-MB-231 cells with new DNA damage (53BP1-mCherry foci) during migration through small (≤ 2 × 5 μm^2^) constrictions (*n* = 381 cells) or 15 × 5 μm^2^ control channels (*n* = 196 cells). *, *p* < 0.05 based on unpaired *t*-test with Welch’s correction. **(F)** Percentage of MDA-MB-231 cells in which new DNA damage during migration through ≤ 2 × 5 μm^2^ constrictions was associated with either NE rupture or with nuclear deformation in the absence of NE rupture. *, *p* < 0.05 based on unpaired t-test with Welch’s correction. **(G)** Association of new DNA damage incurred during migration through ≤ 2 × 5 μm^2^ constrictions with either NE rupture (Rupture) or nuclear deformation without NE rupture (Deformation), for a panel of cell lines. The results correspond to the data presented in Fig. 1C and 1F, and Suppl. Fig. S1G-I. **(H)** Representative image sequence of a MDA-MB-231 cell co-expressing NLS-GFP and 53BP1-mCherry incurring DNA damage during migration in a dense (1.7 mg/ml) collagen matrix. White arrowheads indicate newly occurring 53BP1-mCherry foci. Scale bar: 5 μm **(I)** Percentage of MDA-MB-231 cells with new DNA damage (53BP1-mCherry foci), comparing cells that migrate (*n* = 48 cells) with those that remain stationary (*n* = 29 cells) in a collagen matrix (1.7 mg/ml). *, *p* < 0.0001 based on Fisher’s test. **(J)** Percentage of MDA-MB-231 cells in which new DNA damage during migration through a collagen matrix was associated with either NE rupture or with nuclear deformation in the absence of NE rupture. *n* = 33 cells, *, *p* < 0.05 based on Chi-square test. Data in this figure are presented as mean + S.E.M. See also Figure S1, S2 and S4.

For subsequent studies, we focused on HT1080 and MDA-MB-231 cells to compare and contrast NE rupture and nuclear deformation-associated damage and to identify the underlying mechanism(s). We had previously demonstrated that HT1080 cells migrating through dense collagen matrices exhibit NE rupture and DNA damage [10]. To investigate deformation-induced DNA damage in MDA-MB-231 cells, we imaged cells co-expressing the NLS-GFP and 53BP1-mCherry reporters as they migrated through dense (1.7 mg/ml) collagen matrices using lattice light-sheet microscopy (LLSM). LLSM allows fast, high resolution, 3-D imaging of cells, while minimizing phototoxicity [35]. MDA-MB-231 cells migrated through the collagen matrix with similar velocities as in the microfluidic devices (Suppl. Fig. S1K). Cells migrating through the collagen matrix experienced significantly more DNA damage than cells that remained stationary (Fig. 1H-I). Moreover, this increase in DNA damage was predominantly due to nuclear deformation, independent of NE rupture (Fig. 1H-J, Suppl. Video 3), and the extent of DNA damage increased with the severity of nuclear deformation in individual cells (Suppl. Fig. S1L). These results confirm that MDA-MB-231 cells exhibit deformation-induced DNA damage during migration through confined environments.

### Nuclear compression is sufficient to cause DNA damage

Confined migration involves numerous other cellular processes in addition to nuclear deformation. To test whether nuclear deformation is sufficient to cause DNA damage, we applied external compression to cells cultured on flat, 2-D substrates, resulting in substantial nuclear deformation (Fig. 2A-B). Nuclei were deformed to different heights using a custom-built cell compression device (Fig. 2A) that was inspired by previous designs to study cell confinement and compression [36, 37]. We gradually compressed cells to a height of 2 μm to mimic the nuclear deformation inside the tight constrictions in the microfluidic channels and dense collagen matrices. The cells were compressed for a duration of two hours, similar to their typical transit time through the confined channels in the microfluidic devices (Suppl. Fig. S2A). As baseline control, we used both unconfined cells as well as cells compressed to a height of 5 μm, corresponding to the height of the microfluidic channels. When compressed to 5 μm, the cells deform only moderately (Fig. 2B) compared to unconfined conditions, in which the typical cell height is ≈6-7 μm for MDA-MB-231 cells, but experience similar oxygen and nutrient exchange conditions as in the more severe 2 μm compression case. For some experiments, we additionally tested the effect of intermediate compression (3 μm height).

**Figure 2:**
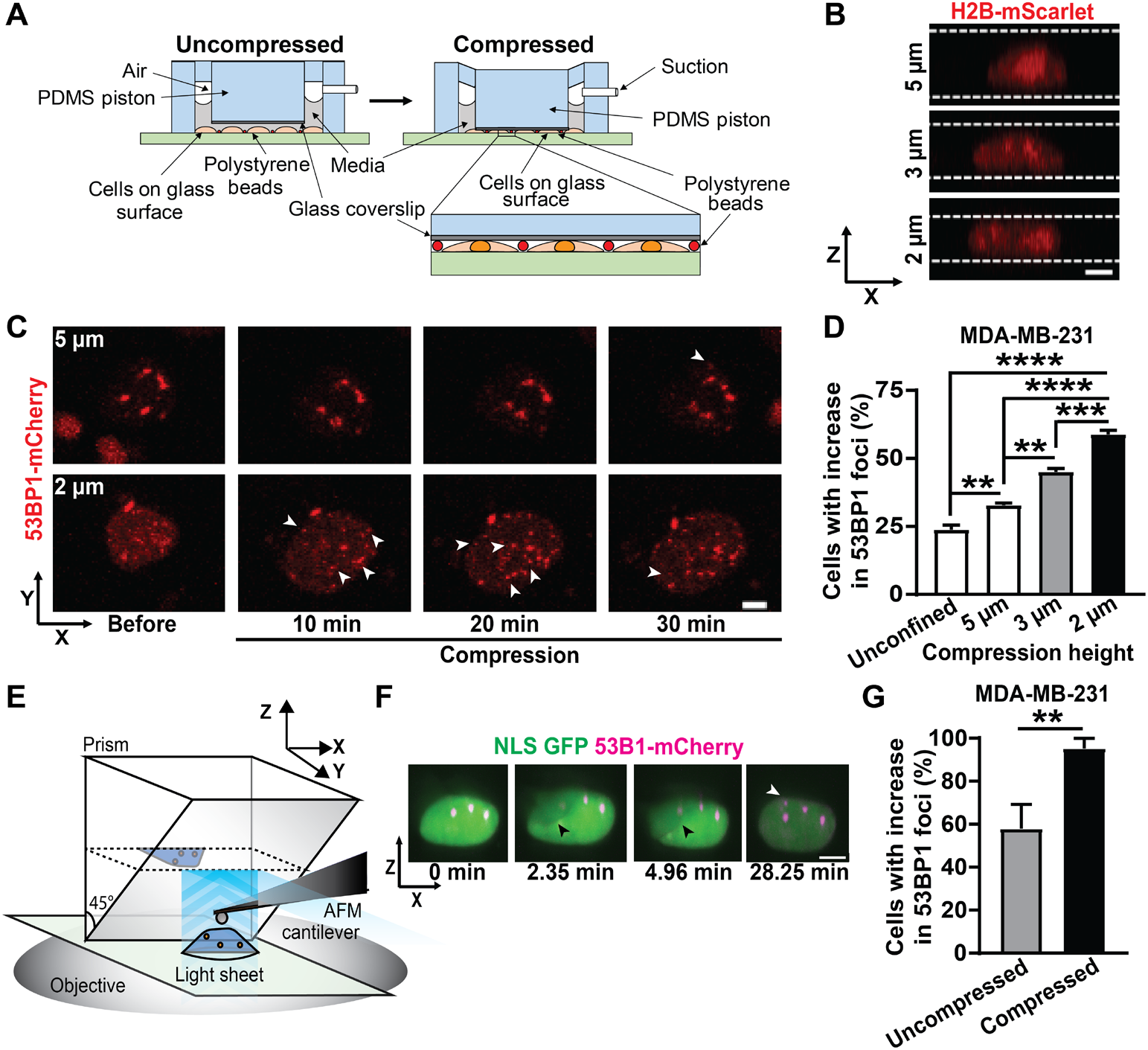
Nuclear compression is sufficient to cause DNA damage. **(A)** Schematic of the custom-built microfluidic compression device with a PDMS piston. The device is connected to a suction source which causes the PDMS piston, with a small circular coverslip attached, to move down onto the cells. Polystyrene beads serve as spacers to ensure a uniform height between the glass coverslip and the glass bottom of the dish. Inset shows cells compressed between the PDMS piston with attached cover slip and the glass surface and the polystyrene beads. **(B)** Representative image sequence showing the nuclear height of a MDA-MB-231 breast cancer cell expressing H2B-mScarlet, compressed to either 5 μm, 3 μm, or 2 μm height using the compression device. White lines indicate the height of the compressed cell. Scale bar: 5 μm **(C)** Representative image sequence showing a MDA-MB-231 breast cancer cell expressing 53BP1-mCherry with new DNA damage formation during compression to either 5 μm or 2 μm (bottom) height. White arrowheads indicate newly occurring 53BP1-mCherry foci; black line indicates the duration of compression. Scale bar: 5 μm **(D)** Percentage of MDA-MB-231 cells with new DNA damage (53BP1-mCherry foci) in unconfined conditions (*n* = 389 cells) or during compression to 5 μm height (*n* = 500 cells), 3 μm height (*n* = 378 cells), or 2 μm height (*n* = 411 cells). **, *p* < 0.01; ***, *p* < 0.001; ****, *p* < 0.0001, based on ordinary one-way ANOVA with Dunnett’s multiple comparison test. **(E)** Schematic overview of the AFM-LS system. A micro-mirror is lowered adjacent to a cell of interest and a vertical light sheet propagates out of the objective illuminating a x-z cross-section of the cell. The image plane is raised to intersect the mirror, capturing the virtual image created by the mirror. The AFM cantilever is positioned between the mirror and cell in order to probe the cell from above while imaging the side-view cross section. **(F)** Representative image sequence showing a MDA-MB-231 breast cancer cell co-expressing NLS-GFP and 53BP1-mCherry experiencing new DNA damage during compression to a height of ~2 μm with an AFM cantilever. Black arrowhead indicates the AFM cantilever; white arrowhead indicates newly occurring 53BP1-mCherry foci. Scale bar: 5 μm **(G)** Percentage of MDA-MB-231 cells with new DNA damage (53BP1-mCherry foci) during compression by an AFM tip (*n* = 21 cells) or in uncompressed control conditions (*n* = 19 cells). **, *p* < 0.01 based on Fisher’s test. Data in this figure are presented as mean + S.E.M. See also Figure S3 and S4.

Compression of the cells increased DNA damage, as visualized by the appearance of new 53BP1-mCherry foci over the entire volume of the nucleus (Fig. 2C). The DNA damage increased with the extent of cell compression, i.e., decreasing nuclear height (Fig. 2C-D). These findings indicate that nuclear deformation is sufficient to cause DNA damage. New DNA damage occurred within 30 minutes of the start of compression (Fig. 2C), thereby ruling out that DNA damage is caused by limited availability of nutrients and oxygen due to compression, which would be expected to result in more gradual increase and later onset of DNA damage [38]. Although compression increased NE rupture (Suppl. Fig. S3A), time-lapse analysis of the NLS-GFP and 53BP1-mCherry reporters revealed that MDA-MB-231 predominantly experienced deformation-induced DNA damage, and not NE rupture-induced DNA damage during compression (Suppl. Fig. S3B), consistent with the results of the confined migration studies (Fig. 1F). Furthermore, while increasing the extent of cell compression from 3 μm height to 2 μm height significantly increased the amount of DNA damage (Fig. 2D), the rate of NE rupture did not increase further with more severe compression (Suppl. Fig. S3A).

To further validate that deformation of the nucleus is sufficient to cause DNA damage, we compressed parts of individual nuclei approximately to a height of 2 μm using an atomic force microscope (AFM) cantilever with a spherical tip (6 μm in diameter) while monitoring nuclear deformation and formation of DNA damage on a light-sheet (LS) microscope (Fig. 2E). The AFM-LS system allows for high-resolution, 3-D imaging of the whole nucleus throughout the compression application [39–41], thereby enabling us to observe the spatio-temporal dynamics of DNA damage (Suppl. Video 4). Experiments in which the spherical tip was brought in contact with the cell without inducing nuclear compression served as controls (Suppl. Video 5). MDA-MB-231 cells showed increased DNA damage upon compression compared to uncompressed controls (Fig. 2F-G). Taken together, these findings support the concept that mechanical deformation of the nucleus or parts of it is sufficient to cause DNA damage.

### Deformation induced DNA damage is independent of reactive oxygen species (ROS)

To investigate the mechanism of deformation-induced DNA damage, we examined the role of ROS, which can increase during confined migration [16] and lead to oxidative DNA damage [42, 43]. However, treatment of MDA-MB-231 with N-acetyl cysteine (NAC), a ROS scavenger [44] that protects cells from H_2_O_2_ induced oxidative stress and apoptosis (Suppl. Fig. S4A) did not reduce DNA damage during confined migration (Suppl. Fig. S4A-B). In contrast, NAC was able to prevent H_2_O_2_ induced apoptosis, serving as a positive control for NAC’s efficacy as a ROS scavenger (Suppl. Fig. S4C). These data suggest that deformation-induced DNA damage in these cells is not caused by increased ROS levels.

### Deformation associated DNA damage occurs specifically in S/G2 phase of the cell cycle

A major cause of DNA damage in proliferating cells is DNA replication stress, which occurs in S/G2 phase of the cell cycle [45]. To test whether DNA damage incurred during confined migration or cell compression was associated with a specific cell cycle stage, we modified MDA-MB-231 and HT1080 cells to stably co-express the 53BP1-mCherry DNA damage reporter and a fluorescent cell cycle reporter, FUCCI (fluorescent ubiquitination-based cell cycle indicator). The FUCCI reporter fluorescently labels cells in the G0/G1 stage of cell cycle in red, and cells in S/G2 phase in green (Fig. 3A; Suppl. Video 6) [46]. We validated that the cell cycle stages determined by the FUCCI reporter were consistent with those obtained by DNA content assay for both MDA-MB-231 and HT1080 cells (Suppl. Fig. S5A-B).

**Figure 3:**
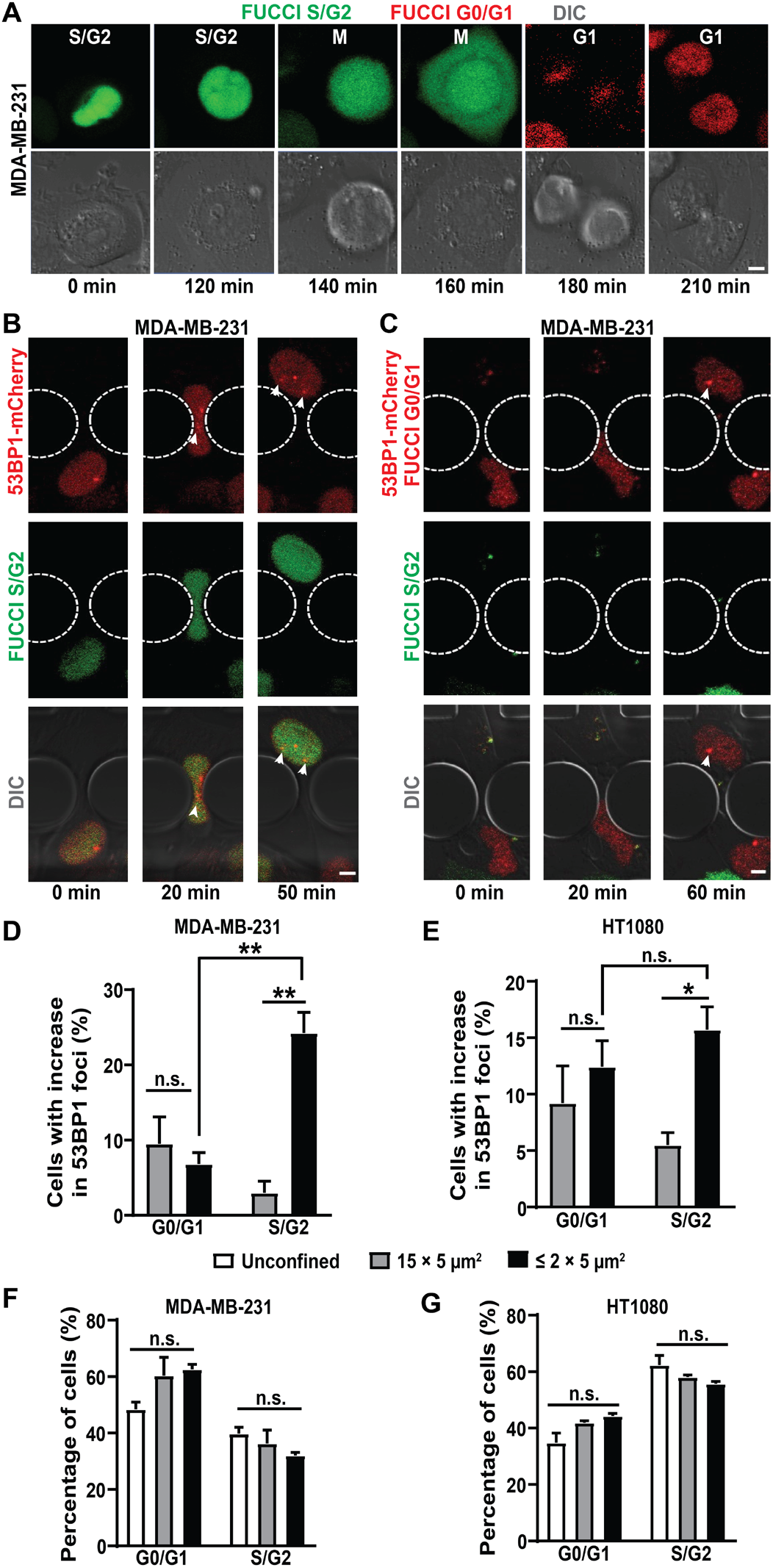
Deformation associated DNA damage occurs specifically in S/G2 phase of the cell cycle. **(A)** Representative image sequence showing a MDA-MB-231 breast cancer cell expressing FUCCI reporter transitioning from S/G2 to M and to G1 cell cycle phase. Scale bar: 5 μm **(B)** Representative image sequence of a MDA-MB-231 breast cancer cell co-expressing FUCCI and 53BP1-mCherry experiencing new DNA damage while in S/G2 phase of the cell cycle during migration through a 2 × 5 μm^2^ constriction. White arrowheads indicate newly occurring 53BP1-mCherry foci. Scale bar: 5 μm **(C)** Representative image sequence of a MDA-MB-231 breast cancer cell co-expressing FUCCI and 53BP1-mCherry experiencing new DNA damage while in G0/G1 phase of the cell cycle during migration through a 2 × 5 μm^2^ constriction. White arrowheads indicate newly formed 53BP1-mCherry foci. Scale bar: 5 μm **(D)** Percentage of MDA-MB-231 cells with new DNA damage (53BP1-mCherry foci) during migration through small (≤ 2 × 5 μm^2^) constrictions (*n* = 327 cells) or 15 × 5 μm^2^ control channels (*n* = 145 cells) as a function of cell cycle phase (G0/G1 or S/G2). **, *p* < 0.01 based on two-way ANOVA with Tukey’s multiple comparison test. **(E)** Percentage of HT1080 fibrosarcoma cells with new DNA damage (53BP1-mCherry foci) during migration through small (≤ 2 × 5 μm^2^) constrictions (*n* = 850 cells) or 15 × 5 μm^2^ control channels (*n* = 371 cells) as a function of cell cycle phase (G0/G1 or S/G2). *, *p* < 0.05 based on two-way ANOVA with Tukey’s multiple comparison test. **(F)** Percentage of MDA-MB-231 cells in G0/G1 or S/G2 phase of the cell cycle in unconfined conditions (*n* = 3544 cells), or during migration through small (≤ 2 × 5 μm^2^) constrictions (*n* = 327 cells) or 15 × 5 μm^2^ control channels (*n* = 145 cells). Differences were not statistically significant (n.s.) based on two-way ANOVA. **(G)** Percentage of HT1080 cells in G0/G1 or S/G2 phase of the cell cycle in unconfined conditions (n = 6108 cells) or during migration through small (≤ 2 × 5 μm^2^) constrictions (*n* = 850 cells) or 15 × 5 μm^2^ control channels (*n* = 371 cells). Differences were not statistically significant based on two-way ANOVA. Data in this figure are presented as mean + S.E.M. See also Figure S5.

Strikingly, in MDA-MB-231 cells, which show predominantly deformation-induced DNA damage, most of the DNA damage during confined migration occurred in S/G2 phase of the cell cycle and not in G0/G1 phase of the cell cycle (Fig. 3B-D). Similar results were obtained for external compression of MDA-MB-231 cells (Suppl. Fig. S5C). In contrast, in HT1080 cells, which predominantly experience NE rupture-induced DNA damage, DNA damage occurred equally in both G0/G1 and S/G2 phase of the cell cycle during confined migration (Fig. 3E) and external compression (Suppl. Fig. S5D). Importantly, confined migration did not select for any particular cell cycle stage, in either MDA-MB-231 cells (Fig. 3F) or HT1080 cells (Fig. 3G). These data suggest that the migration speed and/or efficiency is similar between cells in G0/G1 and S/G2 phase, and the increased occurrence of DNA damage in S/G2 in the MDA-MB-231 cells was not due to an enrichment of cells in this cell cycle phase. The rate of increase in DNA damage was also very similar between cells stably expressing FUCCI and 53BP1-mCherry and those expressing NLS-GFP and 53BP1-mCherry (Suppl. Fig. S5E), suggesting that the increased occurrence of DNA damage in S/G2 in the MDA-MB-231 cells is not due to difficulties in discerning 53BP1-mCherry foci in the G0/G1 phase where cells already express nuclear red fluorescence (Fig. 3C). Moreover, the nuclear deformability for cells in G0/G1 and S/G2 phase of the cell cycle as evaluated using AFM were comparable for both MDA-MB-231 and HT1080 cells (Suppl. Fig. S5F-G), ruling out increased nuclear deformability as the reason behind increased DNA damage in S/G2 phase for MDA-MB-231 cells. Taken together, these findings indicate that NE rupture-induced DNA damage occurs independent of cell cycle stage, consistent with it resulting from the influx of cytoplasmic nucleases, which would attack DNA irrespective of cell cycle stage [21–23] (Fig 6). In contrast, deformation-induced DNA damage occurs primarily in the S/G2 phase of the cell cycle, suggesting that it is linked to DNA replication.

### Deformation-induced DNA damage occurs at replication forks

Since deformation-induced DNA damage occurred primarily in S/G2 phase of the cell cycle, we hypothesized that this damage could be associated with DNA replication stress. DNA replication requires unwinding and subsequent separation of the double strand DNA (dsDNA) into single strands to allow synthesis of the new complementary DNA strands at the replication fork. Replication forks can stall due to conformational and/or torsional stress of the dsDNA, limited availability of nucleotides for DNA synthesis, and other factors [47, 48]. If not repaired in time, stalled replication forks can collapse and form DSBs [49], leading to replication stress.

We stained MDA-MB-231 cells migrating through confined spaces for phosphorylated RPA (p-RPA S33), a marker for single stranded DNA (ssDNA) that accumulates at replication forks, particularly during fork stalling and remodeling [50, 51]. Treatment with hydroxyurea, a DNA replication inhibitor [52–54] and not Phleomycin, a DSB inducing agent [30], led to an increase in number of p-RPA S33 foci, validating its use as a reporter for replication forks and replication stress (Suppl. Fig. S6A). New 53BP1-mCherry foci were frequently co-localized with p-RPA S33 foci (Fig. 4A), suggesting replication stress contributed to the deformation-induced DNA damage during confined migration. To further investigate if this DNA damage occurred at replication forks, we modified MDA-MB-231 and HT1080 cells to co-express 53BP1-mCherry and GFP-PCNA, a fluorescent reporter for DNA replication [55]. PCNA is a DNA clamp that moves along replicating DNA and accumulates at replication forks [56–58]. To validate the GFP-PCNA reporter, HT1080 and MDA-MB-231 cells were treated with hydroxyurea, a DNA replication inhibitor [52–54], which caused a substantial increase in GFP-PCNA foci (Suppl. Fig. S6B-D), indicative of increased replication stress.

**Figure 4:**
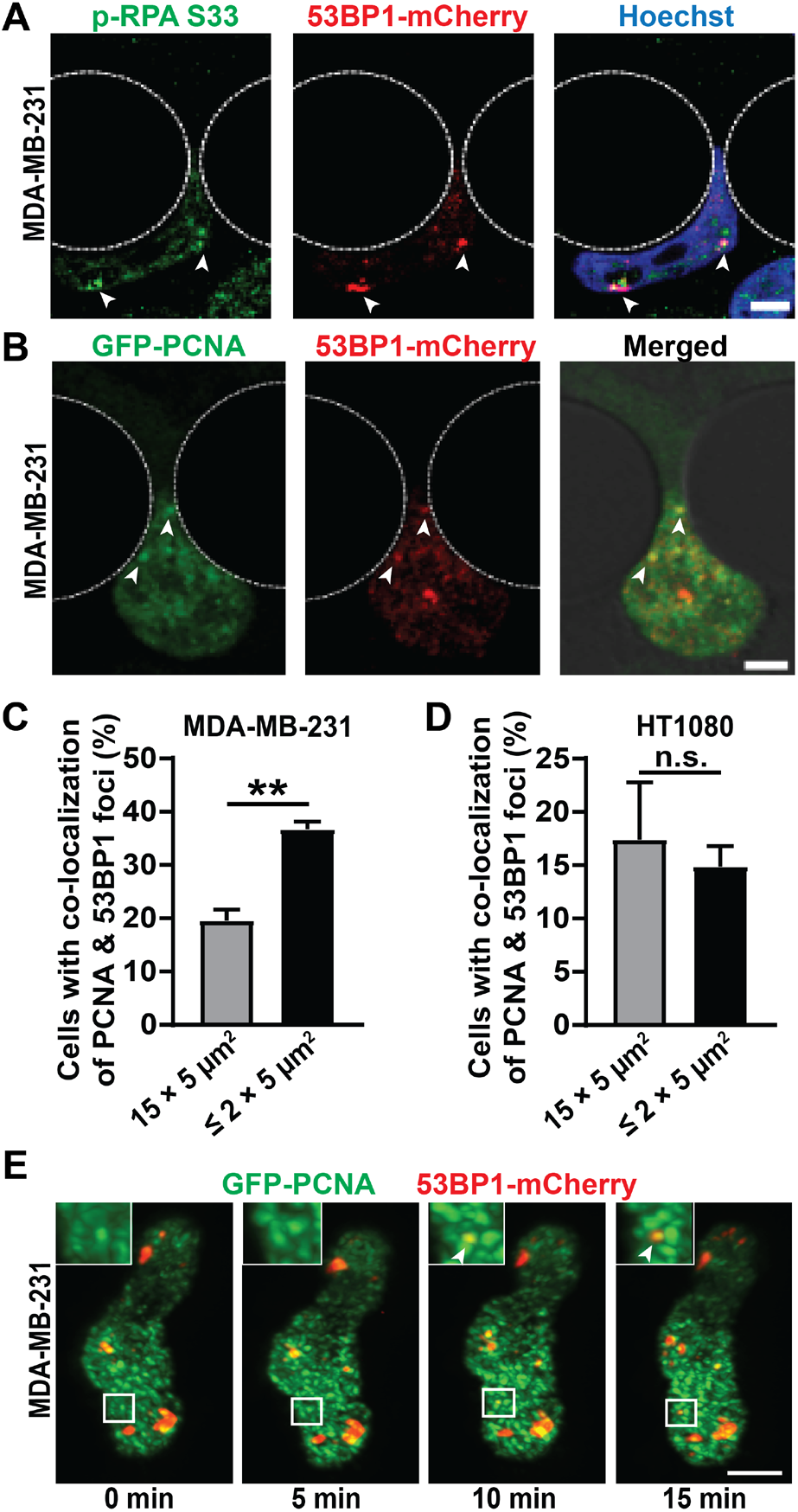
Deformation induced DNA damage occurs at replication forks: **(A)** Representative image panel of a MDA-MB-231 breast cancer cell expressing 53BP1-mCherry during migration through a 1 × 5 μm^2^ constriction, stained for p-RPA S33 to reveal co-localization between p-RPA S33 foci and 53BP1-mCherry foci. White arrowheads indicate sites of co-localization. Scale bar: 5 μm **(B)** Representative image panel showing a MDA-MB-231 breast cancer cell co-expressing GFP-PCNA and 53BP1-mCherry experiencing new DNA damage at replication forks during migration through a 2 × 5 μm^2^ constriction. White arrowheads indicate sites of co-localization between newly occurring 53BP1-mCherry foci and GFP-PCNA foci. Scale bar: 5 μm **(C)** Percentage of MDA-MB-231 cells with co-localization between new DNA damage (53BP1-mCherry foci) and replication forks (GFP-PCNA foci) during migration through small (≤ 2 × 5 μm^2^) constrictions (*n* = 584 cells) or 15 × 5 μm^2^ control channels (*n* = 490 cells). **, *p* < 0.01 based on unpaired *t*-test with Welch’s correction. **(D)** Percentage of HT1080 cells with co-localization between new DNA damage (53BP1-mCherry foci) and replication forks (GFP-PCNA foci) during migration through small (≤ 2 × 5 μm^2^) constrictions (*n* = 986 cells) or 15 × 5 μm^2^ control channels (*n* = 641 cells). Differences were not statistically significant based on unpaired *t*-test. **(E)** Representative image sequence of a MDA-MB-231 cell co-expressing GFP-PCNA and 53BP1-mCherry incurring DNA damage at replication forks during migration in a dense (1.7 mg/ml) collagen matrix. Inset depicts close-up of the region inside the white rectangle to show occurrence of new 53BP1-mCherry foci at replication forks marked by GFP-PCNA foci. White arrowheads indicate site of co-localization between newly occurring 53BP1-mCherry foci and GFP-PCNA foci. Scale bar: 5 μm. Data in this figure are presented as mean + S.E.M. See also Figure S6.

Time-lapse microscopy revealed that new 53BP1-mCherry foci co-localized with GFP-PCNA foci significantly more frequently in MDA-MB-231 cells migrating through the tight constrictions (≤ 2 × 5 μm^2^) compared to cells migrating through the 15 × 5 μm^2^ control channels (Fig. 4B-C; Suppl. Video 7). Similar results were also obtained for BT-549 cells (Suppl. Fig. S6E). In contrast, for HT1080 and RPE-1 cells, which predominantly experience NE rupture-induced DNA damage, co-localization of 53BP1 and PCNA foci was comparable between cells migrating through constrictions and control channels (Fig. 4D; Suppl. Fig. S6F). These findings indicate that in MDA-MB-231 and BT-549 cells, which predominantly exhibit deformation-induced DNA damage, confined migration increases the amount of DNA damage at replication forks compared to baseline levels. MDA-MB-231 cells migrating through collagen matrices showed similar co-localization between new 53BP1-mCherry and GFP-PCNA foci when imaged by LLSM. In these experiments, over 50% of new DNA damage foci appeared at pre-existing PCNA foci, suggesting that the deformation-induced DNA damage occurred at replication forks (Fig. 4E, Suppl. Video 8). Collectively, these findings indicate that deformation-induced DNA damage frequently occurs at replication forks and is likely due to increased replication stress associated with nuclear deformation, which could result in replication fork stalling, collapse, and/or resection.

### Nuclear compression leads to increased replication stress

To directly test whether nuclear deformation can impair DNA replication and cause replication stress, we applied external compression to adherent MDA-MB-231 and HT1080 cells. The compression assay allows precise control of the extent and timing of compression, is suitable for large cell numbers, and enables imaging of live or fixed cells before, during, and after compression. MDA-MB-231 and HT1080 cells were compressed to different heights and analyzed for incorporation of a nucleotide analog, EdU, to assess DNA synthesis rates [59]. MDA-MB-231 but not HT1080 cells subjected to severe compression had significantly reduced EdU incorporation than cells subjected to only mild compression (Fig. 5A-C), suggesting that nuclear deformation reduces DNA replication in MDA-MB-231 cells. To test if the impaired DNA replication during compression in MDA-MB-231 cells might be due to increased replication stress, we stained cells for p-RPA S33 following compression. MDA-MB-231 cells compressed to a height of 2 μm had significantly more p-RPA S33 foci than cells compressed to the 5 μm control height (Fig. 5D-E), indicating that nuclear deformation increases replication stress in these cells, possibly by increasing replication fork stalling, reversal, and/or collapse. In contrast, HT1080 cells did not exhibit any differences in the p-RPA S33 foci upon severe or mild compression (Fig. 5F), suggesting that nuclear deformation impairs DNA replication and increases replication stress only in cells that experience deformation-associated DNA damage and not in cells that experience NE rupture-induced DNA damage.

**Figure 5:**
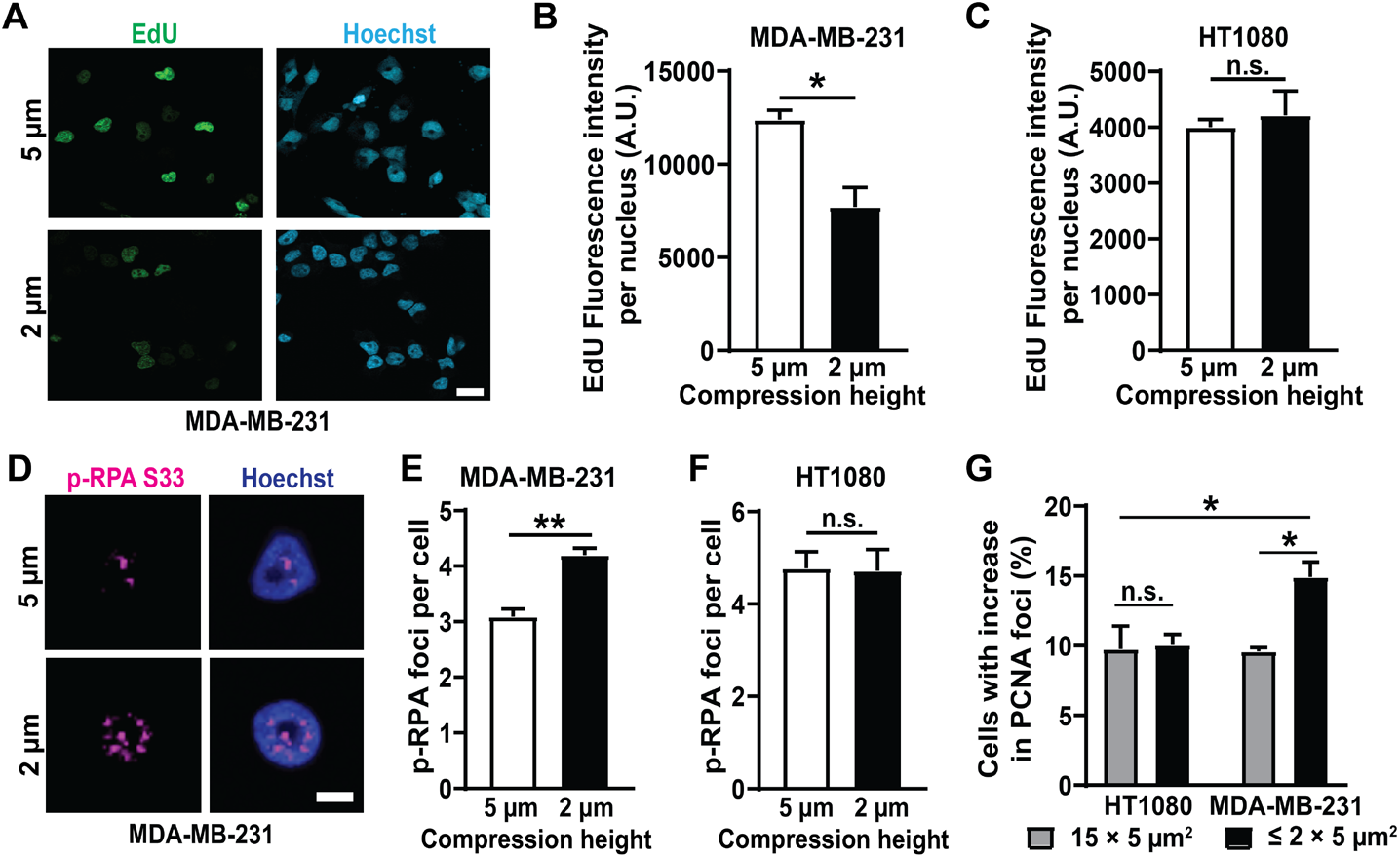
Nuclear deformation leads to increased replication stress. **(A)** Representative images of MDA-MB-231 breast cancer cells showing EdU incorporation during either mild (5 μm height, top) or severe (2 μm height, bottom) compression. Scale bar: 20 μm **(B)** Fluorescence intensity of incorporated EdU per nucleus in MDA-MB-231 cells following compression to either 5 μm height (*n* = 171 cells) or 2 μm height (*n* = 184 cells) for 2 hours. *, *p* < 0.05 based on unpaired *t*-test with Welch’s correction. **(C)** Fluorescence intensity of incorporated EdU per nucleus in HT1080 fibrosarcoma cells following compression to either 5 μm height (*n* = 237 cells) or 2 μm height (*n* = 195 cells) for 2 hours. Differences were not statistically significant based on unpaired *t*-test with Welch’s correction. **(D)** Representative images of a MDA-MB-231 breast cancer cell showing replication forks (p-RPA S33 foci) during compression to either 5 μm or 2 μm (bottom) height. Scale bar: 5 μm **(E)** Average number of p-RPA S33 foci in MDA-MB-231 cells following compression to 5 μm height (*n* = 247 cells) or 2 μm height (*n* = 318 cells) for 2 hours. **, *p* < 0.01 based on unpaired *t*-test with Welch’s correction. **(F)** Average number of p-RPA S33 foci in HT1080 cells following compression to 5 μm height (*n* = 207 cells) or 2 μm height (*n* = 159 cells) for 2 hours. Differences were not statistically significant based on unpaired *t*-test with Welch’s correction. **(G)** Percentage of MDA-MB-231 and HT1080 cells with increase in replication forks (GFP-PCNA foci) during migration through small (≤ 2 × 5 μm^2^) constrictions (*n* = 584 cells for MDA-MB-231; *n* = 986 cells for HT1080) or 15 × 5 μm^2^ control channels (*n* = 490 cells for MDA-MB-231; *n* = 641 cells for HT1080). *, *p* < 0.05 based on two-way ANOVA with Tukey’s multiple comparison test. Data in this figure are presented as mean + S.E.M.

To further investigate whether nuclear deformation during confined migration leads to increased replication stress, we analyzed MDA-MB-231 and HT1080 cells labeled with GFP-PCNA for formation of new PCNA foci as they migrated through the microfluidic device. MDA-MB-231 cells, but not HT1080 cells, exhibited an increase in GFP-PCNA foci as they migrated through the small constrictions (≤ 2 × 5 μm^2^) channels compared to the 15 × 5 μm^2^ control channels (Fig. 5F), indicating increased replication stress in the MDA-MB-231 cells. Taken together, these findings indicate that mechanical deformation of the cell nucleus, for example, during confined migration or external force application, can lead to increased replication stress and DNA damage.

## Discussion

Using a systematic study with a broad panel of cell lines, we showed that DNA damage during confined migration or external cell compression can occur due to at least two separate, but overlapping mechanisms. In some cell lines (MDA-MB-231 and BT-549), nuclear deformation associated with the cell squeezing through tight spaces or external compression is sufficient to cause DNA damage, even without NE rupture, while in other cell lines (HT1080, Human fibroblasts, and RPE-1) DNA damage is predominantly associated with NE rupture. Deformation-induced DNA damage, unlike NE rupture-induced DNA damage, occurs primarily in the S/G2 phase of cell cycle. The deformation-induced DNA damage was located at replication forks, indicating that nuclear deformation is sufficient to cause increased replication stress. We have thus identified a novel mechanism for DNA damage linking mechanical stress on the nucleus and the resulting nuclear deformation to increased replication stress in tumor cells (Fig. 6).

**Figure 6.**
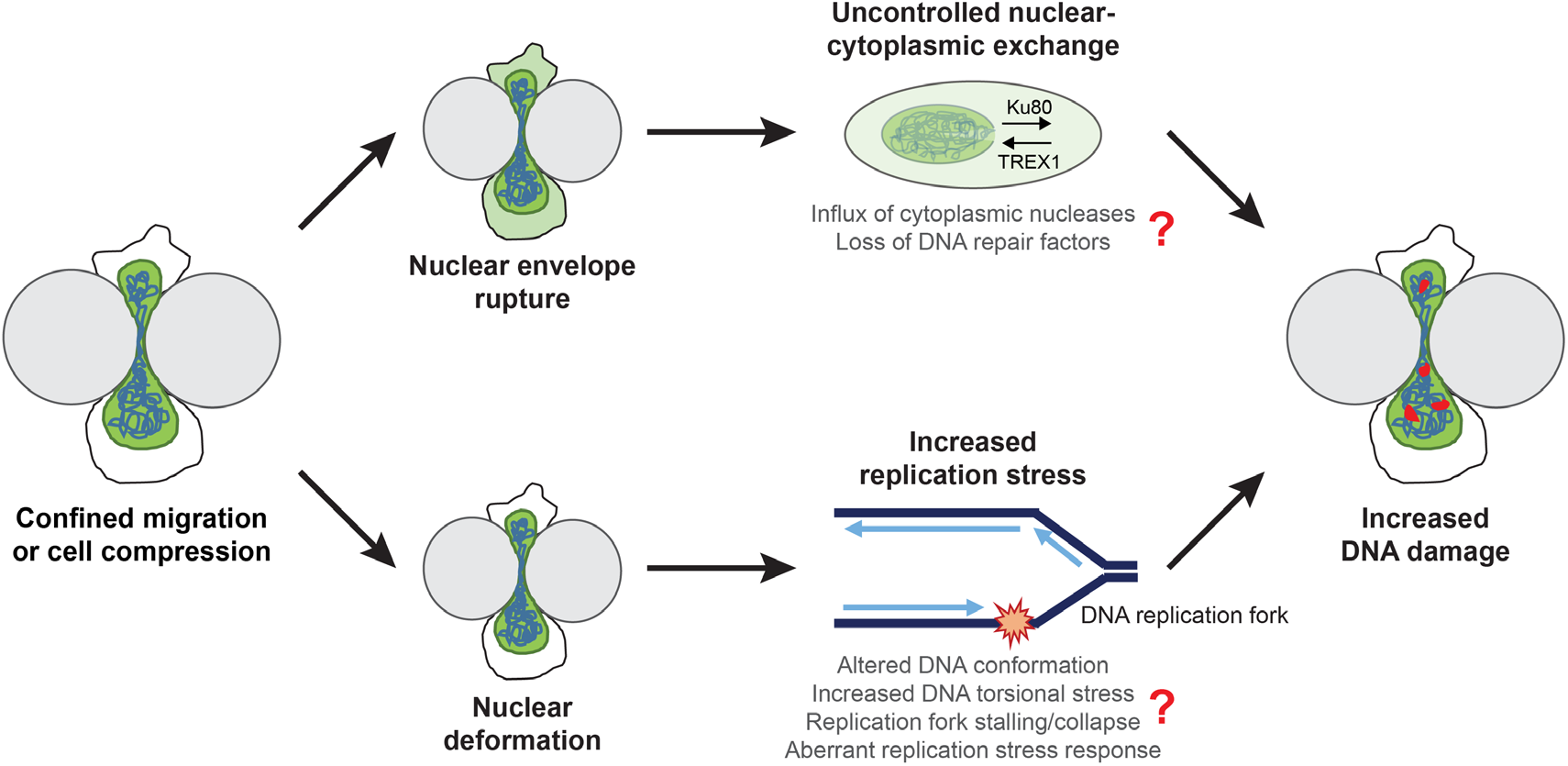
Model for DNA damage during confined migration and cell compression. Nuclei experience severe deformation during cell compression or migration through confined spaces. A subset of cells additionally experience transient NE rupture during these processes. Both nuclear deformation and NE rupture can lead to DNA damage as observed in our experiments, but via separate mechanisms. NE rupture allows uncontrolled exchange of large molecules between the nucleus and cytoplasm. This could lead to a loss of DNA repair factors such as Ku80 or BRCA1 from the nucleus into the cytoplasm, and allow influx of cytoplasmic nucleases such as TREX1 into the nucleus, thereby causing DNA damage. On the other hand, nuclear deformation associated with cell compression or confined migration can alter DNA conformation and make it more difficult to unwind the DNA ahead for replication, leading to an increase in replication stress in the cells. The increased replication stress could be mediated through replication fork stalling or collapse, or aberrant replication stress response by ATR, ultimately resulting in increased DNA damage at replication forks.

While our results indicate that confined migration significantly increases the likelihood of DNA damage, not all cells undergoing confined migration exhibited DNA damage, indicating that it is a stochastic process that may be affected by the extent of nuclear deformation, DNA replication, and other, yet to be identified factors. Since we used a very conservative foci detection approach to avoid false positives, and most of the data were based on imaging only a single focal plane instead of the entire nuclear volume, an approach taken to minimize phototoxicity, our results likely present a lower bound of the confined migration induced DNA damage. Supporting this idea, our experiments using lattice light sheet microscopy to obtain high resolution 3-D images of cells migrating through collagen matrices revealed an increase in DNA damage in close to 80% of all cells (Fig. 1I).

We identified cell line specific differences in the cause of DNA damage during confined migration, however, the molecular mechanisms underlying these differences remain unclear and subject to further investigation. The variability in the susceptibility of different cell lines to deformation-induced DNA damage and NE rupture-induced DNA damage, though not explained by differences in nuclear deformability, migration speed or lamin levels (Suppl. Fig. S2), could arise from multiple other factors, including cell type or tissue of origin. Intriguingly, all of the cell lines tested that exhibited deformation-induced DNA damage had mutations in the gene encoding p53, while the cells exhibiting NE rupture-induced DNA damage had normal p53 function. Thus, it is tempting to speculate that functional p53 provides resistance to deformation-induced replication stress and DNA damage. Consistent with this idea, p53 has been reported to mediate replication stress and restart DNA replication at stalled forks, thereby preventing replication fork collapse and associated DNA damage [60]. At the same time, differences in the susceptibility of cells to NE rupture-induced DNA damage could be due to differences in the expression of cytoplasmic nucleases [21, 23] or other factors.

In contrast to previous reports [15, 16, 27, 61], we did not find evidence of increased cell cycle arrest during confined migration. This could be due to differences in the experimental systems used in the various studies: previous experiments used either transwell plates [15, 16, 27, 61] or long microfluidic channels [15, 16, 27, 61] and assessed cell cycle progression after 12-24 hours of confined migration. In contrast, in our microfluidic devices, only parts of the cell are within the tight constriction at a given time, similar to the situation of cells migrating in collagen matrices [12], and we evaluated cell cycle stage while the cell was moving during the constriction, which typically occurred within 1-3 hours (Fig. 1A). Thus, our results reflect the cell cycle stage at the time new DNA damage occurs, while the findings from previous reports may represent cell cycle arrest resulting from activation of DNA damage response pathways following migration-induced DNA damage.

Our findings suggest that deformation-induced DNA damage occurs at replication forks, which might be stalled or collapsed due to increased replication stress associated with nuclear deformation during confined migration or external compression. The observed increase in p-RPA S33 and GFP-PCNA foci upon nuclear compression and confined migration points towards increase in stalled forks, but we cannot rule out that other mechanisms such as replication fork collapse are responsible for or contribute to the increased DNA damage (Fig. 6). How mechanical forces lead to this increase in replication fork stalling and replication stress, however, remains to be determined. One possible reason could be increased torsional stress in the DNA due to the physical deformation of the nucleus, which could alter DNA conformation and/or make it more difficult to unwind the DNA ahead of the replication fork. Aberrant activity of Mus81, an endonuclease that cleaves stalled replication forks and allows their resolution to prevent DNA DSBs, could also contribute for deformation-induced DNA damage at stalled forks [62]. Moreover, ATR which has important functions in mediating replication stress [45] and has been reported to be mechanosensitive [63], could also play a role in deformation-induced replication stress. Indeed, based on an analysis of the BIOGPS database (www.biogps.org), MDA-MB-231 cells, which experience deformation-induced DNA damage, have lower expression of ATR than HT1080 cells, which primarily exhibit NE rupture-induced DNA damage. This difference may provide one explanation for the increase in replication stress during confined migration in MDA-MB-231 cells.

In conclusion, we uncovered a novel link between mechanically induced nuclear deformation and replication stress. Insights from this work are not only relevant to cells migrating through confining environments, but also to other cells and tissues that experience large compressive forces, for example, during development, in mechanically active tissues such as cartilage, or inside solid tumors [64–66]. Mechanically induced DNA damage could cause cell death, senescence, or, if not repaired properly, lead to mutations, genomic deletions, and/or translocations [14, 67]. Taken together, these findings identify a new mechanism by which mechanical forces can lead to replication stress and increased genomic instability.

## Acknowledgements

The authors thank Connor McGuigan for help with the analysis of cell cycle stage of HT1080 cells in unconfined conditions; Steven Park for helping to optimize the compression device; Jeremy Keys for providing MDA-MB-231 cells expressing H2B-mScarlet; John Heddleston, Eric Wait and the entire HHMI AIC team for help with LLSM image acquisition and processing; E. Timothy O’Brien III for culturing and maintaining cells for the AFM-LS experiments; Robert S. Weiss and Marcus Smolka for DNA damage and replication stress related advice and expertise; and Tyler Kirby, Jeremiah Hsia and other members of the Lammerding laboratory for helpful discussions and support. This work was supported by awards from the National Institutes of Health (R01 HL082792, R01 GM137605, U54 CA210184, and U54 CA193461, to J.L.; NIBIB P41-EB002025 and NSF/NIGMS 1361375 to R.S.; New Innovator DP2 GM229133 and NCI U54 CS21018 to M.J.P), the Department of Defense Breast Cancer Research Program (Breakthrough Award BC150580, to J.L.), the National Science Foundation (CAREER Award CBET-1254846, to J.L. and Award 1752226 to M.J.P), the NSF Graduate Research Fellowship (DGE-1650116 to C.M.H and DGE-1650441 to M.J.C), Caroline H. and Thomas Royster Fellowship (UNC, Chapel Hill to C.M.H) and the Volkswagen Foundation. The AIC is a jointly-funded venture of the Gordon and Betty Moore Foundation and the Howard Hughes Medical Institute. This work was performed in part at the Cornell NanoScale Science and Technology Facility, an NNCI member supported by NSF grant (NNCI-2025233). Image analysis was performed in part through the Cornell University Biotechnology Resource Center, which is supported by New York Stem Cell Foundation (NYSTEM) (CO29155), NIH (S10OD018516), and NSF (1428922).

## Author Contributions

P.S. and J.L. conceptualized and designed all the experiments; C.M.H and R.S. designed AFM experiments; M.J.C and M.J.P designed cell compression device; P.S., C.M.H and S.C. performed experiments; P.S. and J.L. wrote the paper; all authors contributed to the editing of the manuscript; J.L., R.S. and M.J.P acquired funding.

## STAR Methods

### Resource availability

#### Lead Contact

Further information and requests for resources and reagents should be directed to and will be fulfilled by the Lead Contact, Jan Lammerding

#### Materials Availability

This study did not generate any new unique reagents.

#### Data and Code Availability

This study did not generate any new data sets or code.

### Experimental model and subject details

#### Cells and cell culture

The breast adenocarcinoma cell line MDA-MB-231 (ATCC HTB-26) and the breast ductal carcinoma cell line BT-549 (ATCC HTB-122) were purchased from American Type Culture Collection (ATCC); the fibrosarcoma cell line HT1080 (ACC 315) was a gift from Peter Friedl and Katarina Wolf and originally purchased from the DSMZ Braunschweig, Germany; the hTERT-immortalized retinal epithelial cell line RPE-1 was a gift from Marcus Smolka; the SV40-immortalized human fibroblasts were purchased from the Coriell Institute (GM00637). MDA-MB-231, HT1080, RPE-1 and SV40-immortalized human fibroblast cells were cultured in Dulbecco’s Modified Eagle Medium (DMEM) supplemented with 10% (v/v) fetal bovine serum (FBS, Seradigm VWR) and 1% (v/v) penicillin and streptomycin (PenStrep, ThermoFisher Scientific). BT-549 cells were grown in RPMI 1640 media supplemented with 10% FBS and 1% PenStrep. All cell lines were cultured under humidified conditions at 37°C and 5% CO_2_ and verified using STR profiling services from ATCC.

#### Generation of fluorescently labeled cell lines

All cell lines were stably modified with retroviral vectors to express the DNA damage reporter 53BP1-mCherry (mCherry-BP1-2-pLPC-Puro; obtained from Addgene: Plasmid #19835). Some cell lines were additionally modified to stably co-express one or more of the following lentiviral constructs (vectors listed in parenthesis): cell cycle reporter - FUCCI (pLenti6.2-IRES-G1-Orange-S-G2-M-Green-BlastiS, a gift from Katarina Wolf [46]); replication reporter - GFP-PCNA (pHAGE-CMV-GFP-PCNA-BlastiS, a gift from Nima Mosammaparast [55]); nuclear histone marker (pCDH-CMV-H2B-mScarlet-EF1-Puro) and nuclear rupture reporter - NLS-GFP (pCDH-CMV-NLS-copGFP-EF1-blastiS, [10], available through Addgene (#132772)).

### Method details

#### Viral modification

Pseudoviral particles were produced as described previously [68]. In brief, 293-TN cells (System Biosciences, SBI) were co-transfected with the lentiviral plasmid and lentiviral helper plasmids (psPAX2 and pMD2.G, gifts from Didier Trono) using PureFection (SBI), following manufactures protocol. Retroviral particles were produced using 293-GPG cells, which contain the viral packaging plasmid inside. 293-GPG cells were also transfected with plasmid of interest using PureFection. Lentivirus or retrovirus containing supernatants were collected at 48 hours and 72 hours after transfection, and filtered through a 0.45 μm filter. Cells were seeded into 6-well plates so that they reached 50-60% confluency on the day of infection and transduced at most 3 consecutive days with the viral stock in the presence or absence (for BT-549 cell line) of 8 μg/mL polybrene (Sigma-Aldrich). The viral solution was replaced with fresh culture medium, and cells were cultured for 24 hours before selection with 1 μg/mL of puromycin (InvivoGen) or 6ug/mL of blasticidine S (InvivoGen) for 10 days. Cells were also sorted using fluorescence assisted cell sorting (FACS) to ensure expression of all the fluorescent reporters and maintained in a media with antibiotics to ensure continued plasmid expression.

#### Construct validation experiments

The 53BP1-mCherry construct used had a truncated version of the 53BP1 protein containing only the DSB binding domain, preventing downstream signaling [29]. To validate the 53BP1-mCherry construct, cells expressing 53BP1-mCherry were treated with 60 μg/mL Phleomycin (Cayman Chemical), a radiation mimetic agent derived from Streptomyces, which induces DSBs. Cells were imaged every 10-15 min for accumulation of 53BP1-mCherry foci over a period of 1-2 hours after Phleomycin treatment. For GFP-PCNA construct validation, cells expressing GFP-PCNA plasmid were treated with 5 mM hydroxyurea (a gift from Marcus Smolka; ACROS Organics), a DNA replication inhibitor for 24 hours to induce replication fork stalling [53], and increase cellular PCNA expression. Fluorescent images were taken before and after drug treatment, blinded and analyzed for number of PCNA positive cells.

#### Microfluidic migration devices

The devices were prepared as described previously [31, 32]. Microfluidic devices were first assembled by plasma treating the PDMS pieces and coverslips for 5 min, then immediately placing the PDMS pieces on the activated coverslips (pretreated with 0.2M HCl) and gently pressing to form a covalent bond. The finished devices were briefly heated on a hot plate at 95°C to improve adhesion. Devices were filled with 70% ethanol, then rinsed with autoclaved deionized water and coated with extracellular matrix proteins. For all cell lines, except human fibroblasts, devices were coated with 50 μg/mL type-I rat tail collagen (Corning) in acetic acid (0.02 N) overnight at 4°C. For human fibroblasts, devices were incubated with fibronectin (Millipore) in PBS (2-20 μg/mL) overnight at 4°C. After the incubation, devices were rinsed with PBS and medium before loading the cells (about 50,000-80,000 cells per chamber). Subsequently, devices were placed inside a tissue culture incubator for a minimum of 6-24 hours to allow cell attachment before mounting the devices on a microscope for live-cell imaging. The media reservoirs of the device were covered with glass coverslips to minimize evaporation during live cell imaging. Cells were imaged every 5-10 min for 14-16 hours in phenol-red free medium FluoroBrite DMEM supplemented with either 1% or 10% FBS. For experiments with NAC, cells were treated with 10 mM of NAC (Cayman Chemicals) dissolved in PBS starting one hour prior to the start of migration experiments and the treatment was continued for the length of the experiment.

#### Single cell invasion assay in collagen

The single cell invasion assays using collagen matrices were performed as described previously [10]. Briefly, 5 mm round glass coverslips (Warner Instruments) were treated with 1% PEI for 10 min followed by 0.1% Glutaraldehyde for 30 min (to allow consistent bonding between the collagen matrix and the glass coverslip) and washed with PBS before adding the collagen solution. Collagen matrices containing MDA-MB-231 cells were prepared by mixing acidic collagen solution (Corning) (supplemented with DMEM, PBS and NaOH to reach a neutral pH of 7.4) with MDA-MB-231 cells suspended in complete DMEM (density of 100,000 cells /mL). Collagen solution was allowed to polymerize at 37°C for 30 min before adding complete medium. Experiments were carried out 48-72 hours after seeding the cells in the collagen matrix.

#### Cell compression experiments

A custom-built compression device with a PDMS piston (a kind gift from Matthew Paszek) similar to a device published previously [36, 37] and a commercially available version (https://www.4dcell.com/cell-culture-systems/cell-confinement/dynamic-cell-confiner/) was used. Fluorescent beads of either 2, 3 or 5 μm diameter (Invitrogen) were used as spacers to determine the height of compression. Cells were seeded at a density of 10,000/cm^2^ on 35-mm dishes with 14-mm glass coverslips (MatTek) and allowed to adhere overnight inside a tissue culture incubator at 37°C. The next day, cells were washed with PBS, supplemented with FluoroBrite media containing the fluorescent beads, and compressed using the device. To induce suction required to lower the PDMS piston for compression, a syringe pump (New Era Syringe Pump Systems, Inc.) with a 60 mL syringe was operated at a flow rate of 0.5 mL/min for 2 min to achieve compression to a height defined by the spacer beads. Subsequently, the suction was maintained at 0.01-0.1 mL/min to achieve constant compression throughout the course of imaging (2 hours) for all experiments. For release of compression, air was pushed inside at the rate of 0.5 mL/min till the PDMS piston was completely released (about 5-8 min).

#### DNA content analysis

Cell cycle distribution was evaluated using DNA content assay as described previously [69]. Briefly, cells were trypsinized, collected and fixed with 70% Ethanol, on ice for 30 min. Cells were washed and incubated with PBS containing 5 μg/mL RNase A (Omega Bio-Tek) and 10 μg Propidium Iodide (Invitrogen) for 15 min at 37°C, followed by cell sorting using Accuri C6 Flow Cytometer (BD Biosciences). A linear gate was placed on the forward and side scatter plot, to eliminate debris and collect only intact cell data. An additional gate on a secondary graph plotting propidium iodide total fluorescence intensity area vs. height was used to exclude doublets and triplets. Samples were run at 35 μL/min, with a core size of 16 μm, until 100,000 gated cells were recorded. DNA content levels were recorded by plotting propidium iodide intensity vs. cell count on a linear scale. Peaks were used to estimate cell cycle phase, and cell count per phase was determined using the Accuri C6 Plus software (BD Biosciences).

#### Immunofluorescence staining

For validation of the 53BP1-mCherry construct, co-immunofluorescence staining for ɤ-H2AX was performed. Cells cultured on cover slips (pretreated with fibronectin) overnight were fixed with 4% PFA for 20 min at 37°C, permeabilized with PBS containing 0.25% Triton X-100 for 15 min at room temperature, washed, and stained with anti- ɤ-H2AX antibody (Millipore, dilution 1:500). For staining after compression, cells were treated with extraction solution (containing HEPES, NaCl, EDTA, Sucrose, MgCl_2_ and 0.5% Triton X-100) for 15 min on ice followed by fixation with 4% PFA for 20 min at room temperature, permeabilized with PBS containing 0.25% Triton X-100 for 15 min at room temperature, washed, and stained with anti-p-RPA32 (S33) antibody (Bethyl Laboratories, Inc.; dilution 1:1000). For EdU labeling, cells were pulsed with 10 μM EdU (Jena Bioscience) for 2 hours while they were compressed to different heights. After compression, cells were fixed with 4% PFA for 20 min followed by permeabilization with PBS containing 0.25% Triton X-100 for 15 min at room temperature, washed and labelled for EdU using click-chemistry as described previously [59]. For p-RPA staining after Phleomycin or hydroxyurea treatment, cells cultured on cover slips (pretreated with fibronectin) overnight, were treated with either 60 μg/ml of Phleomycin for varying durations or 5 mM of hydroxyurea for 2 hours followed by protein extraction, fixation and staining as mentioned above.

#### Western Blot analysis

Cells were lysed in high salt RIPA buffer containing protease (complete EDTA-Free, Roche) and phosphatase (PhosSTOP, Roche) inhibitors. Protein was quantified using Bio-Rad Protein Assay Dye, and 20–30 μg of protein lysate was separated using a 4–12% Bis-Tris polyacrylamide gel with a standard SDS–PAGE protocol. Protein was transferred to a polyvinylidene fluoride membrane at room temperature at a voltage of 16V for 1 hour. Membranes were blocked using 3% BSA in Tris-buffered saline containing 0.1% Tween-20, and primary antibodies (Lamin A/C (Santa Cruz) - dilution: 1:1000; H3 (Abcam) – dilution: 1:5000) were diluted in the same blocking solution and incubated overnight at 4 °C. Protein bands were detected using IRDye 680LT (LI-COR) secondary antibody, imaged on an Odyssey CLx imaging system (LI-COR) and analysed in Image Studio Lite (LI-COR).

#### Apoptosis assay

Cells were cultured in a 96-well plate at a density of 3000-6000 cells/well and allowed to attach overnight at 37°C Following day, cells were supplemented with Fluorobrite DMEM media and pre-treated with 10 mM NAC for 1 hour at 37°C before being treated with 400 μM H_2_O_2_ for 30 minutes. Cells were analyzed for apoptosis using the NucView 488 Caspase-3 Assay Kit for Live Cells (Biotium) following manufacturer’s protocol. Briefly, cells were treated with 5 μM of the Caspase-3 substrate for 30 minutes at room temperature, before being imaged using the IncuCyte ZOOM (Sartorius) system. All the treatments were continued for the entire length of the experiment.

#### Fluorescence and confocal microscopy

Microfluidic device migration experiments, compression experiments, construct validation experiments, and immunofluorescence staining were imaged on inverted Zeiss Observer Z1 microscopes equipped with temperature-controlled stages (37°C) and CCD camera (Photometrics CoolSNAP KINO) using 20× air (NA = 0.8), 40× water (NA = 1.2) and 63× oil (NA = 1.4) immersion objectives. Airy units for all images were set between 1.5 and 2.5. The image acquisition for migration experiments was automated through ZEN (Zeiss) software with imaging intervals for individual sections between 5-10 min. Images were acquired in a single focal plane, without Z-stacks. For live-cell imaging of cell compression, Z-stacks were acquired from fluorescence, reflection and transmission channels in sequential order every 10 min for 2 hours. Images were also taken before compression was started for direct comparison. Apoptosis assay was imaged on the IncuCyte ZOOM system at 20× once every hour for 24 hours. Four images per well were taken for each time point.

#### AFM-LS

MDA-MB-231 cells were cultured in DMEM/F12 media without phenol red supplemented with 10% FBS (Sigma-Aldrich), 1× antimycotic (Gibco) and 15mM HEPES. A day before the experiment, 50-70% confluent cultures were trypsinized and plated on polyacrylamide gels (stiffness of 55 kPa) coated with collagen as described before [40]. Briefly, 10 μL of activated gel solution was deposited on APTES-treated 40 mm round coverslips, allowed to dry, and attached to a 10 mm diameter glass cloning cylinder (Corning) using vacuum grease (Dow Corning). The gel with cloning rings was treated with EDAC and NHS in PBS, inside the incubator, followed by PBS washes before coating with 50 μg/mL collagen (Rat Tail Type I, Invitrogen) for 30 minutes at 37oC. The collagen solution was washed with PBS and DMEM/F12 medium before adding cells. Cells were plated such that only 1-3 cells were present per field of view at 60× magnification. To compress individual nuclei, we employed the use of atomic force microscopy (AFM) along with light-sheet (LS) microscopy as described previously [39, 40]. Briefly, a beaded cantilever (beaded in house and calibrated using thermal tuning as described in [41]) was mounted onto the AFM and lowered over a cell of interest. A small (180 μm) mirror (Precision Optics Corporation) was placed adjacent to the cell of interest and the objective lens (UplanSAPO 60×/1.2 W, Olympus, Japan) was raised such that the image plane intersected the mirror in order to achieve side-view imaging. A vertical light sheet propagates out of the objective lens and an electrically tunable lens was used to lower the waist of the light sheet such that it lied within the cell. Volumetric images were acquired before, during and after compression. Cells were compressed to an approximate height of 2 μm by lowering the cantilever at the rate of 250 nm/s. For control cells, the cantilever was lowered to indent the cell only a few hundred nanometers. Images were acquired for 20 minutes with compression before retracting the cantilever at the rate of 250 nm/s. Cells were imaged for 5 minutes after retraction as well. The AFM-LS system used has an objective lens heater (Hk-100, Thorlabs, Inc, USA) with a PIV controller (Thorlabs, Inc, USA) and a heated scanning stage (Oxford Instruments, UK) to maintain the temperature of the sample at 37°C.

To study the mechanical properties of nuclei, samples were prepared as described above. A beaded (6 μm diameter) cantilever was aligned directly above the center of the cell nucleus and lowered at a rate of 1 μm/s to compress the nucleus to a height of 2 μm, approximately. Simultaneously, side-view light-sheet images of NLS-GFP were collected at a rate of 4 Hz (200 ms exposure, 50 ms delay) and were used to segment the nucleus during compression. ImageJ was used to extract the nuclear cross-sectional area (NCSA) and the nuclear perimeter (NP). Additionally, the total force response, F, was measured by the AFM with a bandwidth of 2 kHz. The following equation was fit to the indentation portion of the force curve, as described previously [70]

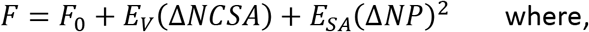

*F*_0_ = any force response prior to deformation of the nucleus;

*E*_*V*_ and *E*_*SA*_ = fitting parameters physically representing the cell’s resistance to bulk and surface deformations of the nucleus, respectively.

*E*_*V*_ is correlated with the nuclear elastic modulus and is primarily dictated by chromatin compaction, while *E*_*SA*_ is correlated with the nuclear stretch modulus and is primarily dictated by the nuclear lamina [70]

#### AFM and Hertzian Analysis

Cells were cultured and prepared as described in the AFM-LS section, above, however, instead of collagen coated polyacrylamide gels, cells were plated directly on collagen-coated coverslips. Cell cycle stage was determined using epifluorescence followed by compression with a beaded (6 μm diameter) cantilever positioned directly on top of the nucleus. Force curves were acquired at an approach velocity of 1 μm/s and data was acquired at 2 kHz bandwidth. Elastic modulus was extracted from the force versus indentation data using the Hertz equation for contact between an elastic sphere and an infinite half-space as described below:

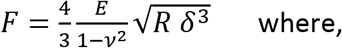

F = force measured by the AFM,

*v* = Poisson ratio (set to 0.5 for this analysis),

R = radius of the indenter,

*δ* = indentation, and

E = elastic modulus.

A custom-written MATLAB program developed by Kellie Beicker (https://cdr.lib.unc.edu/concern/dissertations/d504rk581) was used to extract the elastic modulus from the first 1.5 μm of indentation for each force curve. Contact points were determined algorithmically via a golden-section search method.

#### Lattice light-sheet Microscopy (LLSM)

Single cell invasion assay in collagen was imaged using the lattice light sheet microscope (LLSM) [65] housed in the Advanced Imaging Center at the Howard Hughes Medical Institute Janelia Research Campus. The system is configured and operated as previously described [35]. Samples were illuminated by a 2D optical lattice generated by a spatial light modulator (SLM, Fourth Dimension Displays). The light-sheet pattern was a square lattice with minimum NA of 0.44 and a maximum NA of 0.54. The sample was excited by 488 nm and 560 nm diode lasers (MPB Communications) at 50% AOTF transmittance and 100 mW initial box power through an excitation objective (Special Optics, 0.65 NA, 3.74-mm WD). Fluorescent emission was collected by detection objective (Nikon, CFI Apo LWD 25XW, 1.1 NA), and detected by a sCMOS camera (Hamamatsu Orca Flash 4.0 v2) at 100 ms exposure time. Acquired data were deskewed as previously described [35] and deconvolved using an iterative Richardson-Lucy algorithm. Point-spread functions for deconvolution were experimentally measured using 200nm tetraspeck beads adhered to 5 mm glass coverslips (Invitrogen) for each excitation wavelength.

#### Image analysis

Image sequences were analyzed using ZEN (Zeiss), Zoom (Sartorius) ImageJ [71] or Imaris (BitPlane) using only linear intensity adjustments uniformly applied to the entire image region. For DNA damage analysis, the number of 53BP1-mCherry foci were manually counted in the same cell, before, during, and after each transit through a constriction in the microfluidic channels. Nuclear rupture was detected by an increase of the cytoplasmic NLS-GFP signal. Transit time through the microfluidic devices were calculated using a custom-written MATLAB script [11], available on request. Cell cycle stage was determined by the fluorescence signal of the FUCCI reporter – red was counted as G0/G1 cell cycle stage, and green as S/G2 cell cycle stage. Co-localization of PCNA and 53BP1 was counted manually using image sequences of cells migrating through microfluidic devices. For the compression experiments, DNA damage analysis was performed by manually counting 53BP1-mCherry foci in the maximum intensity projections of confocal image stacks covering the entire nuclear volume. For unconfined conditions, plated cells were monitored for DNA damage over two hours, similar to compression experiments. A total of about 200-1000 cells were counted for each condition in the microfluidic migration and compression experiments. Cells were excluded if there was excessive 53BP1 foci at the start of the experiment preventing analysis of new foci for both migration and compression experiments. For AFM-LS experiments, 53BP1 foci were counted manually by examining the full 3-D image stacks as well as maximum intensity projections. A total of *n* = 21 compressed cells and *n* = 19 control cells were included in the analysis. Three cells (all compression experiments) were excluded from analysis because of excessive 53BP1 foci at the start of the experiment, and two other cells (one control, one compression) were excluded because of ambiguity in whether or not a new 53BP1 focus had formed. For AFM-side view LS experiments to study mechanical properties, two independent experiments were performed with *n* = 17 MDA-MB-231 cells and *n* = 15 HT1080 cells. For collagen experiments, DNA damage analysis was performed by manually counting 53BP1-mCherry foci in the maximum intensity projections of LLSM image stacks covering the entire nuclear volume for both migrating and stationary cells. Nuclear rupture was detected by an increase of the cytoplasmic NLS-GFP signal. A total of *n* = 33 cells were analyzed. Migration speed in the collagen matrices was also calculated using tracked nuclear surfaces over time in Imaris. For DNA damage analysis with respect to degree of nuclear deformation, nuclei were classified into ‘mild’, ‘moderate’ and ‘severe’ deformation, qualitatively and analyzed for new 53BP1-mCherry foci as mentioned above. For PCNA and 53BP1 foci counting in cells migrating through collagen matrices, surfaces were created using Imaris for both PCNA and 53BP1 foci and shortest surface-surface distance was calculated for each tracked point. A total of *n* = 21 cells were analyzed. EdU intensity and p-RPA foci were analyzed using ImageJ. Apoptotic cells were analyzed using Zoom software. An image processing mask was used to calculate percent apoptotic cells after 24 hours of treatment. Graphs were generated in Excel (Microsoft), and figures were assembled in Illustrator (Adobe).

### Quantification and statistical analysis

#### Statistical analysis

Unless otherwise noted, all experimental results are from at least three independent experiments. For data with normal distribution, we used either two-sided Student’s *t*-tests (comparing two groups) or one-way analysis of variance (ANOVA) (for experiments with more than two groups) with post hoc tests. For experiments with two variables and more than two groups, two-way ANOVA was used with post hoc tests. All tests were performed using GraphPad Prism. Welch’s correction for unequal variances was used with all *t*-tests, comparing two groups. One-way ANOVA with post hoc multiple-comparisons testing using Dunnett’s correction was performed to determine differences between 5 μm, 3 μm and 2μm compression heights for all compression experiments involving three different heights, while Tukey’s correction was used for comparison between NAC, control, H_2_O_2_ and combined NAC and H_2_O_2_ groups as well as for comparison between different Phleomycin treated time points, control and hydroxyurea treated groups. For experiments involving individual cells, such as the AFM-LS compression experiments and migration through collagen matrices, a Chi-square or Fisher’s test was used. Post hoc multiple-comparisons testing with Tukey’s correction was used with all two-way ANOVA analyses, comparing two variables and more than two groups. Statistical details of each experiment can be found in the figure legend. Unless otherwise indicated, error bars represent s.e.m.

## Supplementary Videos

**Supplementary Video 1: HT1080 cell migrating through a 2 × 5 μm^2^ constriction in the microfluidic device. Related to Figure 1.** Representative example of HT1080 cell co-expressing NLS-GFP (green) and 53BP1-mCherry (red) migrating through a small constriction (2 × 5 μm^2^) in the microfluidic device. The video shows NE rupture, as evidenced by the increase in cytoplasmic NLS-GFP (denoted by red arrow), followed by new DNA damage, as seen by occurrence of 53BP1-mCherry foci. White arrowheads indicate newly occurring 53BP1-mCherry foci. Scale bar: 5 μm

**Supplementary Video 2: MDA-MB-231 cell migrating through a 2 × 5 μm^2^ constriction in the microfluidic device. Related to Figure 1.** Representative example of MDA-MB-231 cell co-expressing NLS-GFP (green) and 53BP1-mCherry (red) migrating through a 2 × 5 μm^2^ constriction, undergoing severe nuclear deformation and exhibiting new DNA damage as seen by the increase in 53BP1-mCherry foci. White arrowheads indicate newly occurring 53BP1-mCherry foci. Scale bar: 5 μm

**Supplementary Video 3: MDA-MB-231 cell migrating through collagen matrix**. Related to **Figure 1.** Representative example of MDA-MB-231 cell co-expressing NLS-GFP (green) and 53BP1-mCherry (red) migrating inside a 1.7 mg/ml collagen matrix. The cell exhibits severe nuclear deformation and DNA damage, as seen by the formation of 53BP1-mCherry foci. White arrowheads indicate newly occurring 53BP1-mCherry foci. Scale bar: 10 μm

**Supplementary Video 4: MDA-MB-231 cell undergoing AFM compression. Related to Figure 2.** Representative example of MDA-MB-231 cell co-expressing NLS-GFP (green) and 53BP1-mCherry (red) subjected to external compression to a height of 2μm with an AFM tip. DNA damage formation can be seen by occurrence of new 53BP1-mCherry foci, indicated by white arrows. Scale bar: 5 μm

**Supplementary Video 5: MDA-MB-231 ‘control’ cell for AFM compression experiments. Related to Figure 2.** Representative example of MDA-MB-231 cell co-expressing NLS-GFP (green) and 53BP1-mCherry (red) in which the AFM tip is only brought in gentle contact with the cell surface without causing nuclear deformation. In this case, no new DNA damage is formed, and only pre-existing 53BP1-mCHerry foci are visible. Scale bar: 5 μm

**Supplementary Video 6: MDA-MB-231 cell expressing FUCCI. Related to Figure 3.** Representative example of MDA-MB-231 cell co-expressing FUCCI cell cycle reporter and 53BP1-mCherry (red foci). The fluorescence signal of the FUCCI reporter switches from green to red as the cell transitions from S/G2 (green) to M (green) and G0/G1 (red). Scale bar: 5 μm

**Supplementary Video 7: MDA-MB-231 cell exhibiting DNA damage at replication forks in microfluidic device. Related to Figure 4.** Representative example of MDA-MB-231 cell co-expressing GFP-PCNA (green) and 53BP1-mCherry (red) migrating through a small (≤ 2 × 5 μm^2^) microfluidic constriction. The cell experiences DNA damage, as seen by the formation of new 53BP1-mCherry foci, which co-localize with sites of stalled replication forks, visibly by the GFP-PCNA foci. White arrowheads indicate sites of new 53BP1 foci and PCNA co-localization. Scale bar: 5 μm

**Supplementary Video 8: MDA-MB-231 cell exhibiting DNA damage at replication forks in collagen matrix. Related to Figure 4.** Representative example of MDA-MB-231 cell co-expressing GFP-PCNA (green) and 53BP1-mCherry (red) migrating through a 1.7 mg/ml collagen matrix. The cell exhibits nuclear deformation and DNA damage, seen by the formation of new 53BP1-mCherry foci, which co-localize with sites of existing GFP-PCNA foci. Scale bar: 5 μm

