## Supplementary Figures and Figure Legends for "Nuclear deformation causes DNA damage by increasing replication stress"

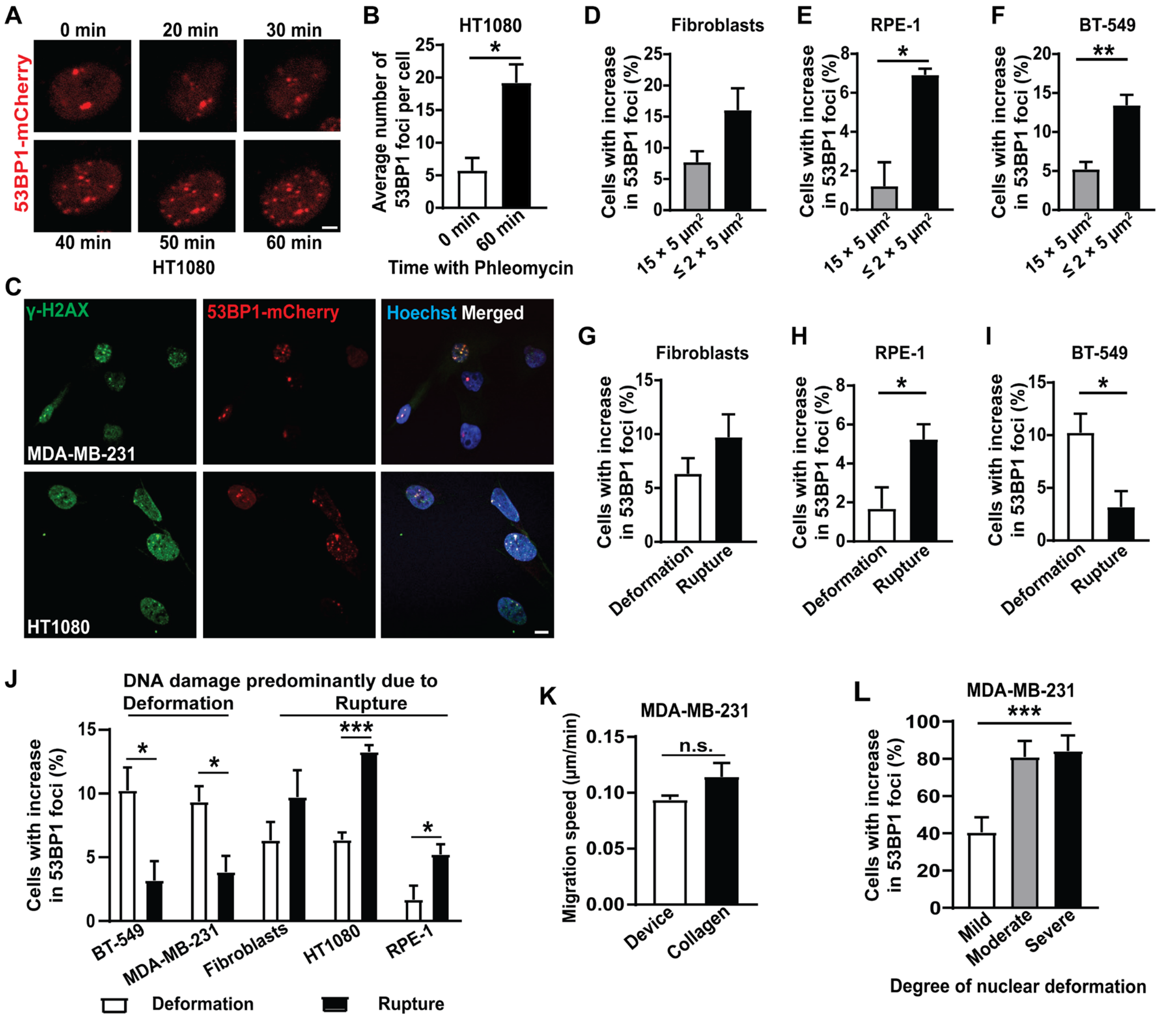


**Supplementary Figure S1: 53BP1-mCherry construct validation and DNA damage due to migration through confined spaces. Related to Figure 1. (A)** Representative image sequence of a HT1080 fibrosarcoma cell expressing 53BP1-mCherry treated with Phleomycin, showing increasing number of 53BP1-mCherry foci over time, indicating increasing DNA damage. Scale bar: 5 µm **(B)** Percentage of HT1080 cells with new DNA damage (53BP1-mCherry foci) before Phleomycin treatment and after 60 minutes of Phleomycin treatment. $n$ = 86 cells; *, $p < 0.05$ based on unpaired $t$-test with Welch's correction. **(C)** Representative image panel showing co-localization between ɣ-H2AX foci and 53BP1-mCherry foci in MDA-MB-231 and HT1080 cells. Scale bar: 10 µm **(D)** Percentage of human fibroblast cells with new DNA damage (53BP1-mCherry foci) during migration through small (≤ 2 × 5 µm$^2$) constrictions ($n$ = 230 cells) or 15 × 5 µm$^2$ control channels ($n$ = 135 cells). **(E)** Percentage of RPE-1 retinal epithelial cells with new DNA damage (53BP1-mCherry foci) during migration through small (≤ 2 × 5 µm$^2$) constrictions ($n$ = 407 cells) or 15 × 5 μm$^2$ control channels (*n* = 259 cells). *, $p < 0.05$ based on unpaired *t*-test with Welch's correction. **(F)** Percentage of BT-549 breast cancer cells with new DNA damage (53BP1-mCherry foci) during migration through small (≤ 2 × 5 μm$^2$) constrictions (*n* = 555 cells) or 15 × 5 μm$^2$ control channels (*n* = 260 cells). **, $p < 0.01$ based on unpaired *t*-test with Welch's correction. **(G)** Percentage of human fibroblasts cells in which new DNA damage during migration through ≤ 2 × 5 μm$^2$ constrictions was associated with either NE rupture or with nuclear deformation in the absence of NE rupture. *n* = 230 cells **(H)** Percentage of RPE-1 cells in which new DNA damage during migration through ≤ 2 × 5 μm$^2$ constrictions was associated with either NE rupture or with nuclear deformation in the absence of NE rupture. *n* = 407 cells; *, $p < 0.05$ based on unpaired *t*-test with Welch's correction. **(I)** Percentage of BT-549 cells in which new DNA damage during migration through ≤ 2 × 5 μm$^2$ constrictions was associated with either NE rupture or with nuclear deformation in the absence of NE rupture. *n* = 555 cells; *, $p < 0.05$ based on unpaired *t*-test with Welch's correction. **(J)** Percentage of cells in which new DNA damage during migration through ≤ 2 × 5 μm$^2$ constrictions was associated with either NE rupture (Rupture) or with nuclear deformation in the absence of NE rupture (Deformation), for a panel of cell lines. The results correspond to the data presented in Fig. 1C and 1F, and Suppl. Fig. S1G-I. *, $p < 0.05$; ***, $p < 0.001$ based on unpaired *t*-test with Welch's correction. **(K)** Migration speed of MDA-MB-231 cells during migration through ≤ 2 × 5 μm$^2$ constrictions in the microfluidic device (*n* = 537 cells) and collagen matrices (1.7 mg/ml of collagen; *n* = 27 cells). Differences were not statistically significant based on Chi-square test. **(L)** Percentage of MDA-MB-231 breast cancer cells with new DNA damage (53BP1-mCherry foci) due to mild (*n* = 37 cells), moderate (*n* = 21 cells), or severe (*n* = 19 cells) nuclear deformation during migration through a collagen matrix (1.7 mg/ml). ***, $p < 0.001$ based on Chi-square test. Data in this figure are presented as mean + S.E.M.

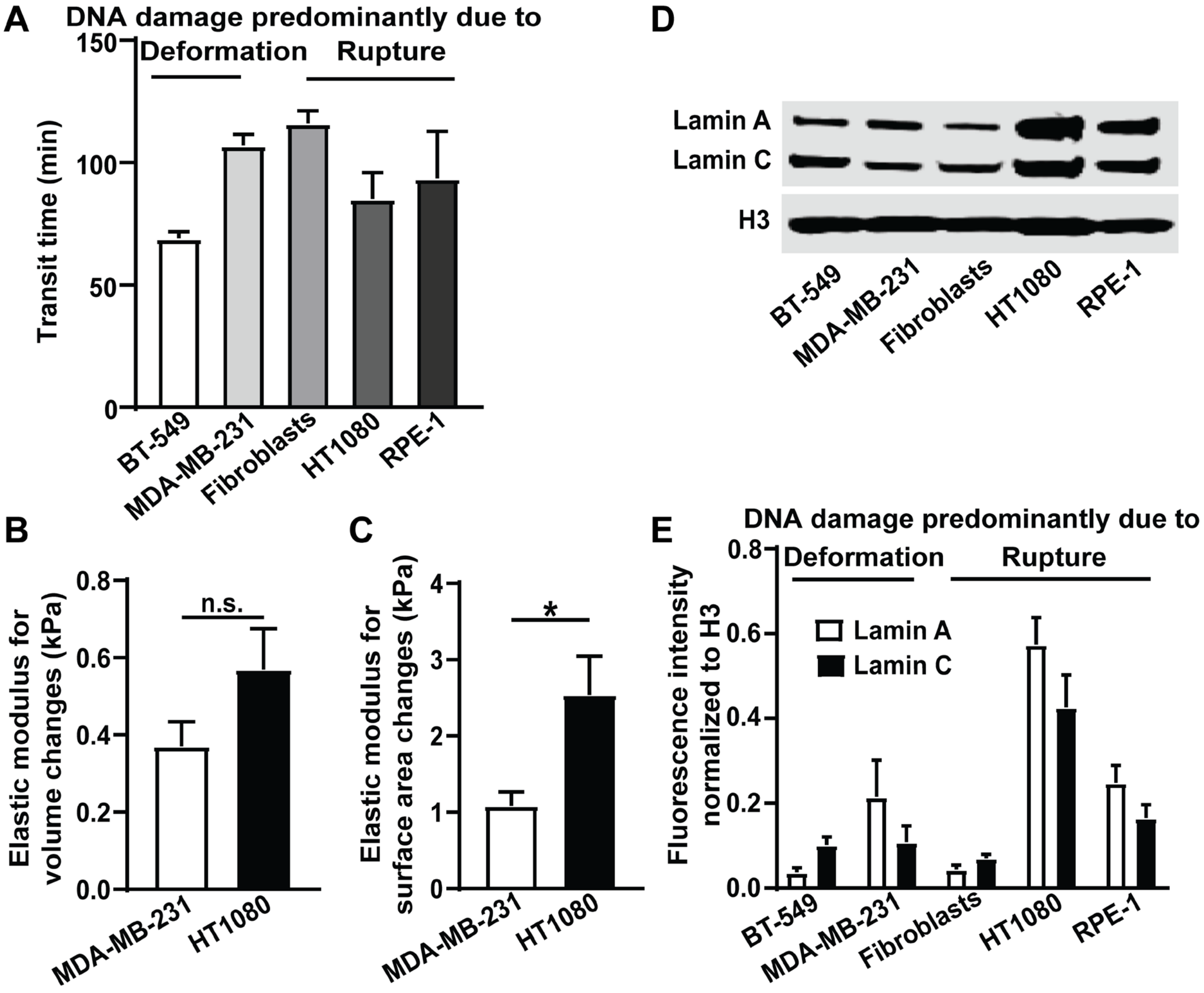


**Supplementary Figure S2: Transit time, nuclear deformability, and lamin A/C level analysis in a panel of cell lines. Related to Figure 1. (A)** Transit time for BT-549 (*n* = 444 cells), MDA-MB-231 (*n* = 381 cells), human fibroblasts (*n* = 125 cells), HT1080 (*n* = 326 cells), and RPE-1 (*n* = 269 cells) cells to migrate through ≤ 2 × 5 $\mu m^2$ constrictions in the microfluidic device. **(B)** Elastic modulus for MDA-MB-231 (*n* = 17 cells) and HT1080 (*n* = 15 cells) cells for bulk nuclear deformation by a beaded AFM tip. Differences were not statistically significant based on unpaired *t*-test with Welch's correction. **(C)** Elastic modulus for nuclear surface area deformation for MDA-MB-231 (*n* = 17 cells) and HT1080 (*n* = 15 cells) cells probed with a beaded AFM tip. *, $p < 0.05$ based on unpaired *t*-test with Welch's correction. **(D)** Representative western blot of lamin A and C levels in a panel of human cell lines. Histone H3 was used as a loading control. **(E)** Quantification of lamin A and C levels based on *N* = 3 western blot experiments. Data in this figure are presented as mean + S.E.M.

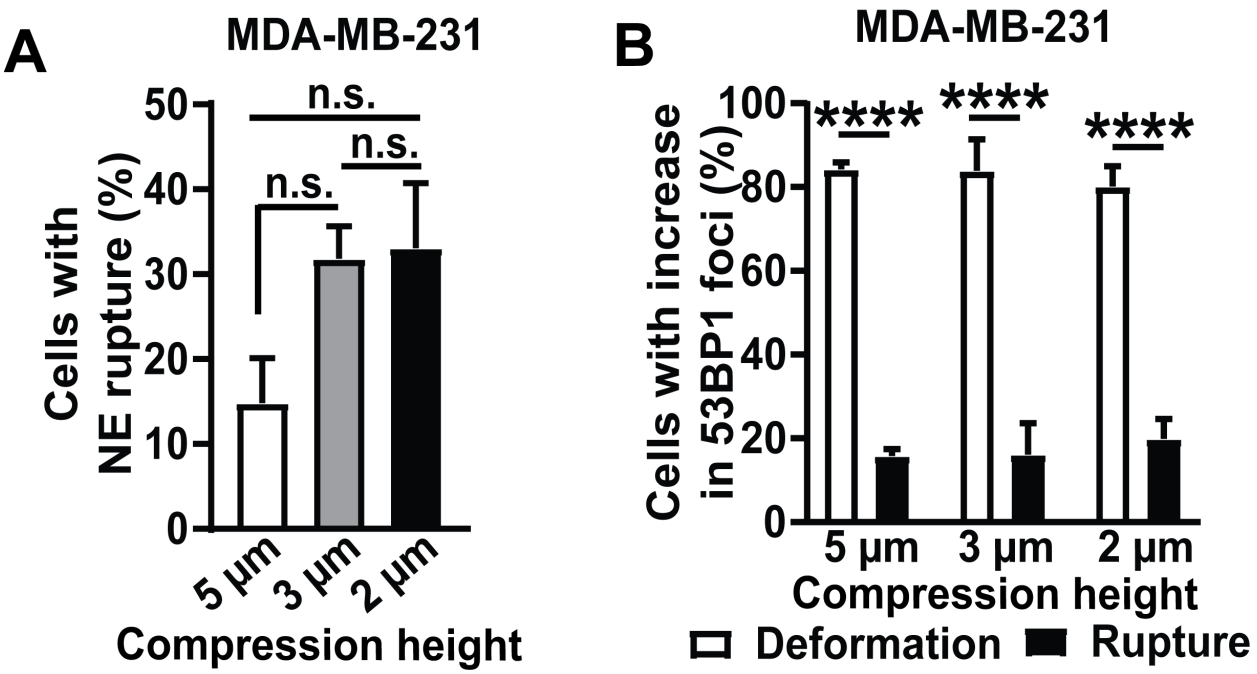


**Supplementary Figure S3: Cell compression causes NE rupture and DNA damage. Related to Figure 2. (A)** Percentage of MDA-MB-231 cells that experience NE rupture during external compression to 5 µm height (*n* = 500 cells), 3 µm height (*n* = 378 cells), or 2 µm height (*n* = 411 cells). Differences were not statistically significant based on one-way ANOVA with Dunnett's multiple comparison test. **(B)** Percentage of MDA-MB-231 cells in which new DNA damage during compression to 5 µm, 3 µm, or 2 µm height was associated with either NE rupture or with nuclear deformation without NE rupture. ****, $p < 0.0001$ based on two-way ANOVA with Tukey's multiple comparison test. Data in this figure are presented as mean + S.E.M.

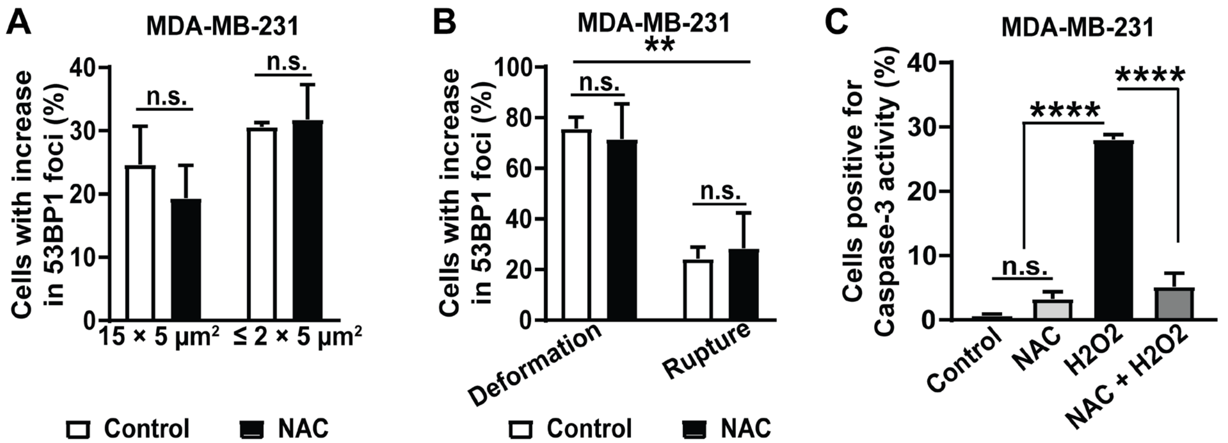


**Supplementary Figure S4: NAC treatment does not rescue DNA damage. Related to Figure 1,2.** **(A)** Percentage of MDA-MB-231 cells treated with or without NAC that exhibit new DNA damage (53BP1-mCherry foci) during migration through small (≤ 2 × 5 μm$^2$) constrictions (*n* = 181 cells for vehicle control; *n* = 182 cells for NAC treatment) or 15 × 5 μm$^2$ control channels (*n* = 95 cells for vehicle control; *n* = 83 for NAC treatment). Differences were not statistically significant based on two-way ANOVA with Tukey's multiple comparison test. **(B)** Percentage of MDA-MB-231 cells treated with or without NAC in which new DNA damage during migration through small (≤ 2 × 5 μm$^2$) constrictions was associated with either NE rupture or with nuclear deformation in the absence of NE rupture. **, $p < 0.01$ based on two-way ANOVA with Tukey's multiple comparison test. **(C)** Percentage of MDA-MB-231 cells undergoing apoptosis (positive for Caspase-3 activity) after 24 hours of NAC, $H_2O_2$ and combined NAC and $H_2O_2$ treatment. N = 3 experiments. ****, $p < 0.0001$ based on one-way ANOVA with Tukey's multiple comparison test. Data in this figure are presented as mean + S.E.M.

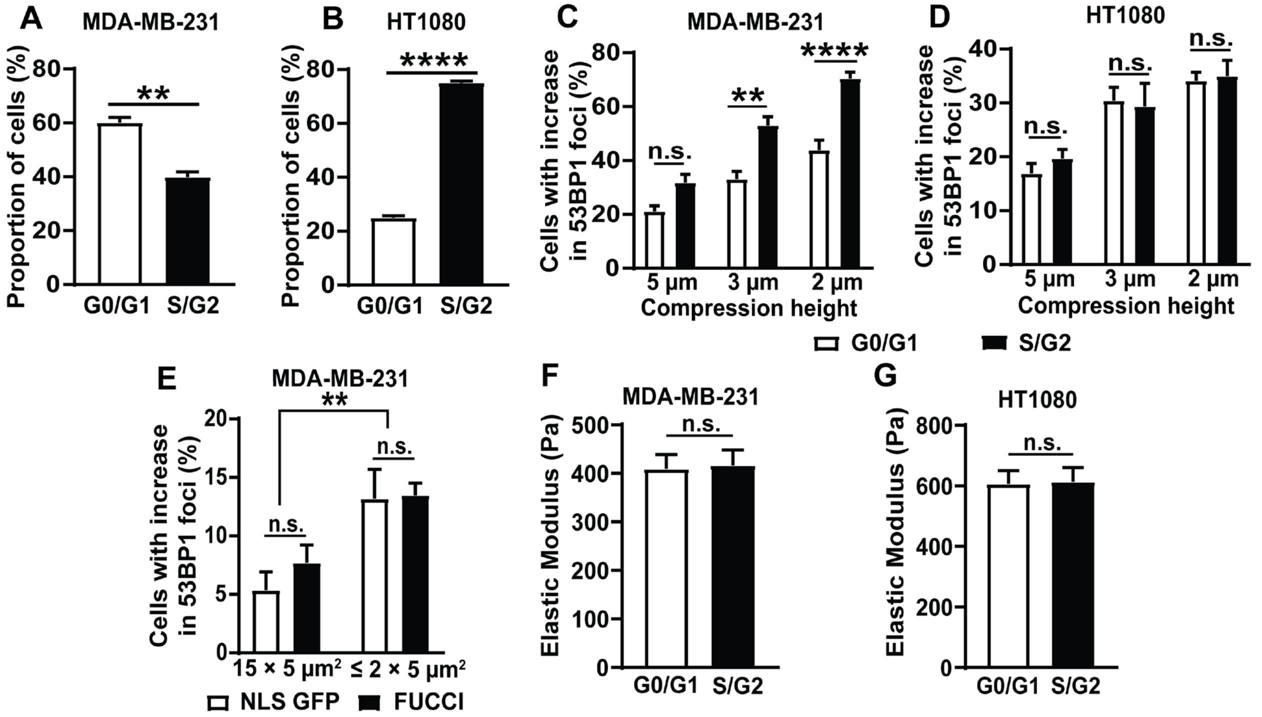


**Supplementary Figure S5: Cell cycle stage analysis by DNA content assay and during nuclear compression. Related to Figure 3. (A)** Proportion of MDA-MB-231 cells in G0/G1 phase or S/G2 phase of the cell cycle, determined by DNA content assay. **, $p < 0.01$ based on unpaired *t*-test with Welch's correction. **(B)** Proportion of HT1080 cells in G0/G1 phase or S/G2 phase of the cell cycle, determined by DNA content assay. ****, $p < 0.0001$ based on unpaired *t*-test with Welch's correction. **(C)** Percentage of MDA-MB-231 cells with new DNA damage (53BP1-mCherry foci) in G0/G1 or S/G2 phase of the cell cycle during compression to either 5 µm ($n = 574$ cells), 3 µm ($n = 359$ cells), or 2 µm ($n = 522$ cells) height. **, $p < 0.01$; ****, $p < 0.0001$ based on two-way ANOVA with Tukey's multiple comparison test. **(D)** Percentage of HT1080 cells with new DNA damage (53BP1-mCherry foci) in G0/G1 or S/G2 phase of the cell cycle during compression to either 5 µm ($n = 656$ cells), 3 µm ($n = 449$ cells) or 2 µm ($n = 672$ cells) height. Differences were not statistically significant based on two-way ANOVA. **(E)** Percentage of MDA-MB-231 cells expressing NLS-GFP or FUCCI reporter with new DNA damage (53BP1-mCherry foci) during migration through small ≤ 2 × 5 µm$^2$ constrictions ($n = 381$ for NLS-GFP cells and $n = 327$ for FUCCI cells) or 15 × 5 µm$^2$ control channels ($n = 196$ for NLS-GFP cells and $n = 145$ for FUCCI cells). **, $p < 0.01$ based on two-way ANOVA with Tukey's multiple comparison test. **(F)** Nuclear elastic modulus, measured by AFM indentation with a spherical tip probe, for MDA-MB-231 cells in G0/G1 ($n = 50$ cells) or S/G2 ($n = 50$ cells) phase of cell cycle. Cell cycle phase was determined based on the FUCCI reporter expressed in these cells. Differences were not statistically significant based on unpaired *t*-test with Welch's correction. **(G)** Nuclear bulk elastic modulus for HT1080 cells in G0/G1 ($n = 51$ cells) or S/G2 ($n = 51$ cells) phase of cell cycle. Differences were not statistically significant based on unpaired *t*-test with Welch's correction. Data in this figure are presented as mean + S.E.M.

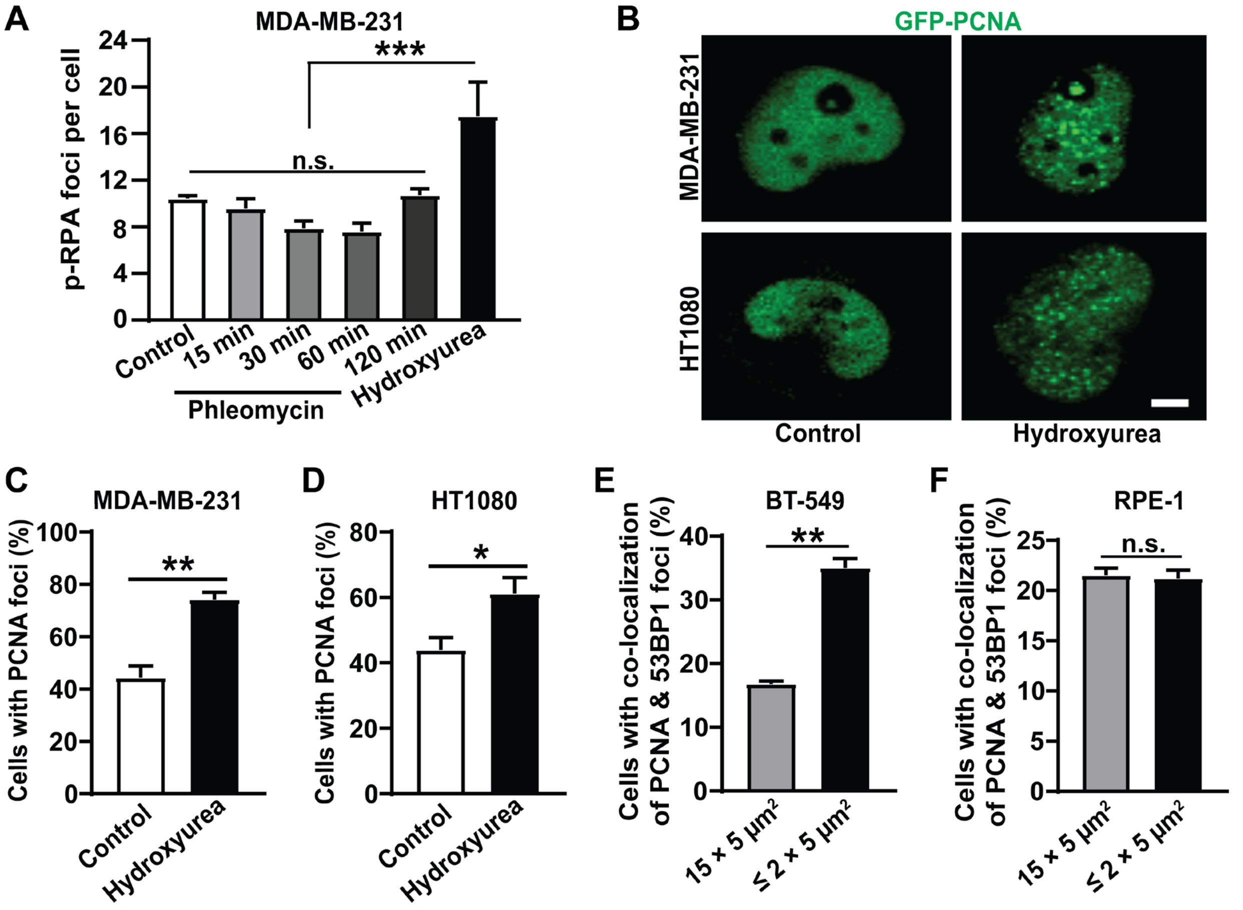


**Supplementary Figure S6: GFP-PCNA construct validation and replication stress experiments. Related to Figure 4. (A)** Average number of p-RPA S33 foci in MDA-MB-231 untreated control cells ($n$ = 235 cells) or following treatment with Phleomycin for 15 minutes ($n$ = 201 cells), 30 minutes ($n$ = 195 cells), 60 minutes ($n$ = 187 cells), or 120 minutes ($n$ = 191 cells). Hydroxyurea treatment (120 minutes) was used as a positive control for replication stress ($n$ = 158 cells). ***, $p < 0.001$ based on one-way ANOVA with Tukey's multiple comparison test. **(B)** Representative image sequence of MDA-MB-231 and HT1080 cells expressing GFP-PCNA, showing an increase in GFP-PCNA foci, indicating stalled replication forks, following treatment with hydroxyurea. Scale bar: 5 µm. **(C)** Percentage of MDA-MB-231 cells with GFP-PCNA foci following hydroxyurea treatment or vehicle control. $n$ = 147 cells for control; $n$ = 163 cells for hydroxyurea treatment; **, $p < 0.01$ based on unpaired $t$-test with Welch's correction. **(D)** Percentage of MDA-MB-231 cells with GFP-PCNA foci following hydroxyurea treatment or vehicle control. $n$ = 132 cells for control; $n$ = 143 cells for hydroxyurea treatement; *, $p < 0.05$ based on unpaired $t$-test with Welch's correction. **(E)** Percentage of BT-549 breast cancer cells with co-localization between new DNA damage (53BP1-mCherry foci) and replication forks (GFP-PCNA foci) during migration through small (≤ 2 × 5 µm$^2$) constrictions ($n$ = 154 cells) or 15 × 5 µm$^2$ control channels ($n$ = 106 cells). **, $p < 0.01$ based on unpaired $t$-test with Welch's correction. **(F)** Percentage of RPE-1 retinal epithelial cells with co-localization between new DNA damage (53BP1-mCherry foci) and replication forks (GFP-PCNA foci) during migration through small (≤ $2 \times 5$ $\mu m^2$) constrictions ($n$ = 249 cells) or $15 \times 5$ $\mu m^2$ control channels ($n$ = 126 cells). Differences were not statistically significant based on unpaired $t$-test. Data in this figure are presented as mean + S.E.M.
